# Autoimmune Alleles at the Major Histocompatibility Locus Modify Melanoma Susceptibility

**DOI:** 10.1101/2021.08.12.456166

**Authors:** James Talwar, David Laub, Meghana Pagadala, Andrea Castro, McKenna Lewis, Georg E. Luebeck, Bryan Gorman, Cuiping Pan, Frederick N. Dong, Kyriacos Markianos, Richard Hauger, Saiju Pyarajan, Philip S. Tsao, Gerald P. Morris, Rany M. Salem, Wesley K. Thompson, Kit Curtius, Maurizio Zanetti, Hannah Carter

## Abstract

Autoimmunity and cancer represent two different aspects of immune dysfunction. Autoimmunity is characterized by breakdowns in immune self-tolerance, while impaired immune surveillance can allow for tumorigenesis. The class I major histocompatibility complex (MHC-I), which displays derivatives of the cellular peptidome for immune surveillance by CD8+ T cells, serves as a common genetic link between these conditions. As melanoma-specific CD8+ T-cells have been shown to target melanocyte-specific peptide antigens more often than melanoma-specific antigens, we investigated whether vitiligo and psoriasis predisposing MHC-I alleles conferred a melanoma protective effect. In individuals with cutaneous melanoma from both The Cancer Genome Atlas (N = 451) and an independent validation cohort (N = 586), MHC-I autoimmune allele carrier status was significantly associated with a later age of melanoma diagnosis. Furthermore, MHC-I autoimmune allele carriers were significantly associated with decreased risk of developing melanoma in the Million Veterans Program cohort (OR = 0.962, p = 0.024). Existing melanoma polygenic risk scores (PRS) did not predict autoimmune allele carrier status, suggesting these alleles provide new risk-relevant information. Mechanisms of autoimmune protection were neither associated with improved melanoma-driver mutation association nor improved gene-level conserved antigen presentation relative to common alleles (population frequency ≥ 1%). However, autoimmune alleles showed higher affinity relative to common alleles for particular windows of melanocyte conserved antigens suggesting a potential relationship between antigen processing, binding, and cell-surface presentation. Overall, this study presents evidence that MHC-I autoimmune risk alleles modulate melanoma risk unaccounted for by current PRS.

## Introduction

The incidence of cutaneous melanoma, the most common form, has seen an increase globally, particularly in Western countries.^1,2^ Early detection is a major determinant of overall disease prognosis with the 5-year survival rate dropping precipitously from 99% to 27.3% for local versus distant disease, respectively.^3^ Models developed for early disease detection are often built around well-known environmental and host risk factors including ultraviolet radiation exposure,^4–6^ pigmentary phenotypes,^7–9^ melanocytic nevi count,^10,11^ sex,^12,13^ age,^12,13^ telomere length,^14,15^ immunosuppression,^16,17^ and family history.^18,19^ However, though cutaneous melanoma ranks among the most heritable forms of cancer with an estimated heritability of 58%,^20^ the majority of genetic susceptibility remains unaccounted for.

Cutaneous melanoma is also considered among the most immunogenic forms of cancer. Melanoma exhibits one of the highest mutation burdens across cancers, which is driven primarily by the mutagenic influence of ultraviolet radiation exposure.^21,22^ This increases the number of neoepitopes presented to the immune system and plays an essential role in immune surveillance. Immunosuppression, though, impairs the immune system’s cytotoxic potential and is a documented risk factor for increased melanoma incidence.^16,17^ Lymphocyte infiltration and melanoma-specific antibodies have been shown to be powerful prognostic factors as well.^23,24^ Immune traits themselves show considerable heritability,^25–27^ and early investigations suggest that heritable immune alleles also contribute to melanoma risk.^28–30^ In contrast to cancer, where poor immune function is a risk factor,^31,32^ increased sensitivity of the immune system can lead to autoimmune disorders.^33^ This dichotomy has led to speculation that induction of autoimmunity in cancer patients could lead to tumor regression and better immunotherapy efficacy,^34–39^ though studies investigating the relationship between autoimmunity and cancer risk have returned mixed findings.^40–42^ If autoimmune alleles can enhance host anti-tumor immune responses when immunotherapy is administered, it is possible that they could also enhance anti-tumor immunity more generally. Thus, we speculated that autoimmune alleles, particularly those related to T cell responses directed against melanocyte or melanoma-specific antigens, might modify polygenic risk for melanoma in ways that might not be captured by current polygenic risk scores (PRS).

The class I major histocompatibility complex (MHC-I) represents a fundamental component of the antigen-directed immune response common to cancer and autoimmunity. MHC-I binds and displays peptide antigens derived primarily from intracellular proteins on the cell surface for immune surveillance by CD8+ T-cells.^43^ In cancer, neopeptides containing somatic mutations unique to the tumor genome, when displayed by MHC-I, can be recognized as foreign by CD8+ T cells, triggering the release of cytotoxic granules.^44,45^ The peptide-binding specificity of MHC-I is determined by three highly polymorphic genes, *HLA-A*, *HLA-B* and *HLA-C*, encoded at the Human Leukocyte Antigen (HLA) locus on chromosome 6. The specific set of HLA alleles carried by an individual has been found to impose selective constraints on the developing tumor genome ^46–48^ and modify response to immunotherapy.^49–51^ MHC-I loss, often due to deletion or mutation of HLA genes, is one mechanism of immune evasion during tumor development.^52,53^

MHC-I also plays a role in several skin-specific autoimmune disorders. In vitiligo, destruction of melanocytes and consequent loss of skin pigmentation is mediated by CD8+ T-cell responses to self-antigens displayed by MHC-I.^54,55^ Another skin autoimmune disorder involving CD8+ T cell responses is psoriasis,^56,57^ which is characterized by dermal leukocyte infiltration and hyperproliferation of keratinocytes.^57^ CD8+ T-cells in psoriasis have been shown to target melanocytes in patients carrying particular MHC-I alleles.^58,59^ Both of these conditions also share a risk variant rs9468925 in HLA-B/HLA-C,^60^ supporting a shared MHC-I driven etiology, which is consistent with disease co-occurrence findings in clinical investigations.^61,62^

Intriguingly, multiple vitiligo risk alleles implicated by genome-wide association studies have been shown to exhibit protection from cutaneous melanoma,^28,29^ and emergence of vitiligo during immunotherapy treatment of melanoma patients has been associated with better responses.^63–66^ Though psoriasis has also been documented as an immune-related adverse effect of immunotherapy, the association with melanoma prognosis is far less characterized.^67,68^

Altogether these findings support the potential for class I autoimmune HLA alleles to modify melanoma risk and inform risk prediction. To further investigate this possibility, we evaluated autoimmune HLA carrier status in cutaneous melanoma samples from The Cancer Genome Atlas (TCGA). Skin autoimmune alleles were associated with a significantly later age at diagnosis among melanoma cases, which was recapitulated in a validation cohort of 586 individuals assembled from the UK Biobank and 4 other published melanoma genome sequencing studies. We further investigated the peptide-specificity of autoimmune alleles for a set of 215 melanoma-specific driver mutations spanning 172 genes, and for conserved and cancer antigens previously implicated in melanoma-directed immunity. These analyses highlight a protective role for MHC-I in the context of melanoma development.

## Results

### MHC-I Autoimmune Alleles Associate with a Later Age of Diagnosis in Melanoma

To investigate the relationship between autoimmunity and melanoma development, we sought evidence that MHC-I autoimmune alleles could provide protection from melanoma. We defined a set of 8 MHC-I alleles based on documented links to skin autoimmune conditions. This set included 3 alleles linked to psoriasis (HLA-B*27:05, HLA-B*57:01, HLA-C*12:03),^69–73^ 2 alleles linked to vitiligo (HLA-A*02:01, HLA-B*13:02),^74–79^ and 1 allele linked to both conditions (HLA-C*06:02).^78–81^ We also included 2 alleles (HLA-B*39:06, HLA-B*51:01) with postulated psoriasis associations,^69,82^ but strong associations with other autoimmune conditions, specifically type 1 diabetes ^83–85^ and Behcet’s disease,^86^ respectively.

Using individuals with skin cutaneous melanoma tumors (SKCM) from the TCGA as our discovery cohort (**Supplementary Table 1**), we called MHC-I genotypes using two exome based methods: POLYSOLVER and HLA-HD (*Methods: Datasets*).^87,88^ Autoimmune allele frequencies in the discovery cohort were present at the population distributions as reported by the National Marrow Donor Program (NMDP)^89^ (**Supplementary Table 2**). Individuals under 20 years of age were also excluded from further analysis given their increased likelihood of harboring rare germline predisposing risk variants.

To evaluate the effect of carrying an autoimmune (AI) allele, we partitioned the discovery cohort into two groups: those with at least one AI allele and those lacking any of these alleles. We first assessed the potential for AI allele carrier status to be confounded by sex or UV exposure, two factors that influence melanoma incidence. Men have a well-documented higher risk of developing melanoma,^90,91^ however, we found no significant sex specific age differences across AI allele status (**Supplementary Fig. 1A**, p = 0.247). UV-associated mutational signatures correlated strongly with overall tumor mutation burden in the discovery cohort (Pearson R = 0.984; **Supplementary Fig. 1D**). There were also no significant differences in mutation burden or UV-associated mutational signatures between AI allele carriers and non-carriers (**Supplementary Fig. 1B-D**; A*02:01 included: p_mutation_ = 0.365, p_UV_ = 0.186; A*02:01 excluded: p_mutation_ = 0.352, p_UV_ = 0.415).

We hypothesized that in an all-cancer cohort, a protective effect would manifest as delayed disease onset relative to individuals without these alleles. As HLA-A*02:01 had a high frequency relative to other alleles, and thus could potentially dominate the analysis, we considered AI carrier status both including and excluding HLA-A*02:01. Across the discovery cohort, AI carrier status including HLA-A*02:01 failed to significantly associate with age (**Supplementary Fig. 1E**, p = 0.316). However, when HLA-A*02:01 was excluded, AI allele carrier status was significantly associated with a median later age of melanoma diagnosis of 5 years (**Fig. 1A**, p = 0.002). For subsequent analyses we therefore defined AI allele carrier status based on the other 7 AI alleles; individuals carrying only HLA-A*02:01 were considered non-carriers.

**Figure 1:**
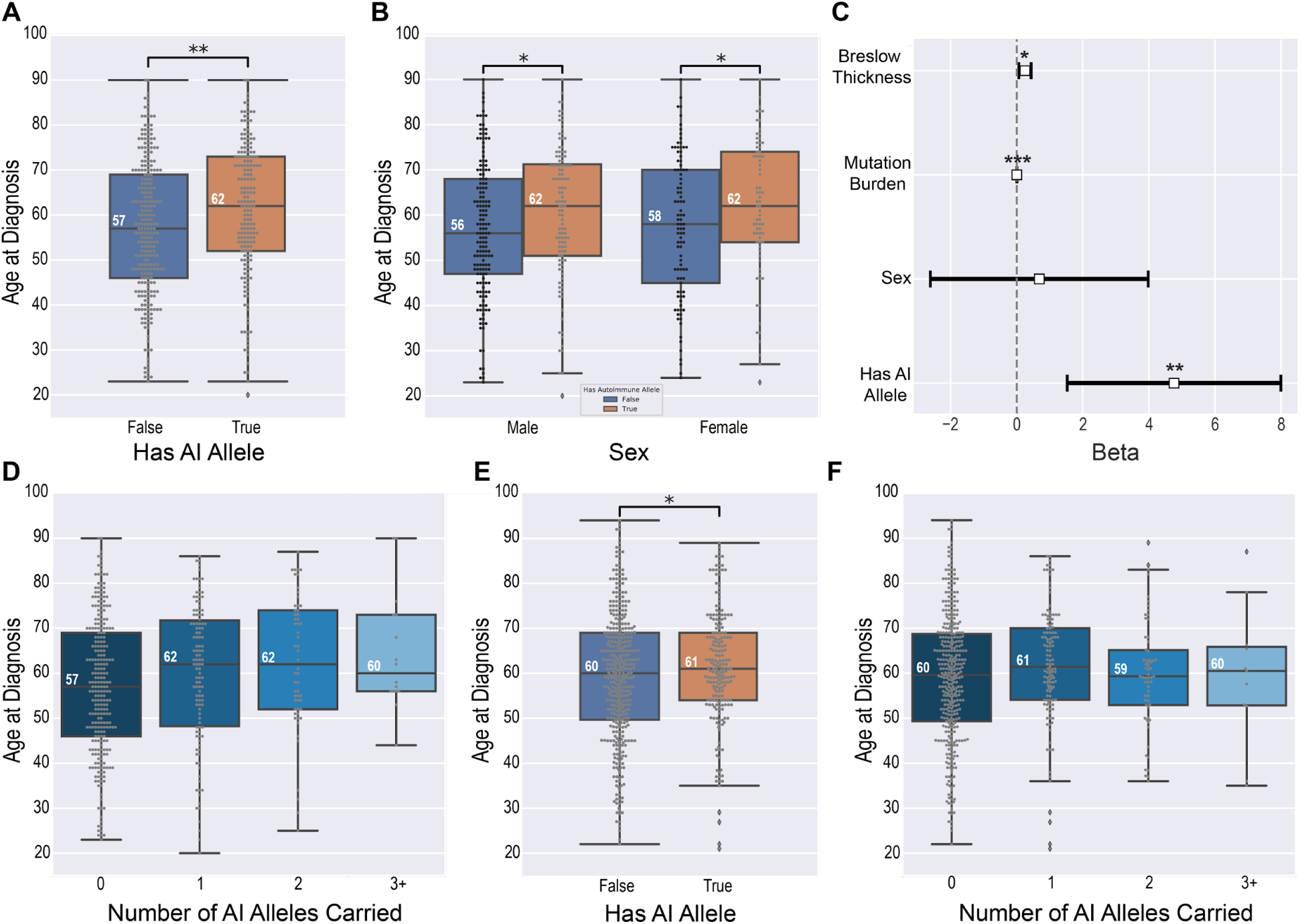
Effect of MHC-I autoimmune alleles on age at diagnosis in melanoma. A) *TCGA*: Having at least one MHC-I linked autoimmune allele is associated with a significant median later age of melanoma diagnosis of 5 years (p = 0.002). B) *TCGA*: MHC-I linked autoimmune allele age of melanoma diagnosis effect is conserved across sex (p_male_ = 0.011, p_female_= 0.033). C) *TCGA:* Having at least one MHC-I linked autoimmune allele is significantly associated with 4.76 delayed years to melanoma diagnosis after controlling for tumor thickness, sex, and mutation burden (p_autoimmune_ = 0.003). Mutation burden also remained significant, but had a minimal contribution to age of diagnosis with 0.004 delayed years until melanoma diagnosis (p_mutation_ = 0.002) D) *TCGA:* Age of diagnosis significantly increases with an individual’s total number of autoimmune MHC-I autoimmune alleles (p = 0.002). E) *Validation*: Having at least one MHC-I links autoimmune allele is associated with a significant median later age of melanoma diagnosis of 1.0 years (p = 0.0435). F) *Validation:* Age of diagnosis increases with an individual’s total number of autoimmune MHC-I autoimmune alleles (p = 0.328).

For the revised definition of carrier status, later age of diagnosis was conserved across sex with a median age difference of 6 years in males and 4 years in females (**Fig. 1B**, p_male_ = 0.011, p_female_= 0.033). We used an ordinary least squares (OLS) regression to model the effect of carrying an AI allele on age at diagnosis with tumor thickness, sex, and mutation burden as covariates and these findings remained significant with a predicted 4.76 delayed years to melanoma diagnosis (**Fig. 1C**, p = 0.003). As tumor thickness was only available for a subset of the discovery cohort (N = 347), we also evaluated the effect of AI allele carrier status with primary vs. metastatic disease, sex, and mutation burden as covariates (N = 416), and sex and mutation burden only as covariates (N = 451). Both cases yielded results consistent with our previous findings, with predicted 3.62 (**Supplementary Fig. 2A**, p = 0.015) and 4.07 (**Supplementary Fig. 2B**, p = 0.005) delayed years to melanoma diagnosis respectively. To ensure no single AI allele drove these findings we conducted a leave-one-out analysis, evaluating AI age effects across all 7 single allele exceptions. Regardless of which allele was held out, we consistently observed a significant relationship between AI status and age of diagnosis (**Supplementary Fig. 3**), with a 5 year later median age of melanoma diagnosis (median age without AI = 57; median age with AI = 62). We did not observe significant differences at the level of HLA supertype, suggesting that the effects are specific to individual autoimmune alleles (**Supplementary Fig. 4**), though the B44 supertype, to which none of the AI alleles belong, trended towards an earlier age of diagnosis (p = 0.199, median earlier age of diagnosis difference = 4 years). We further evaluated whether carrying multiple AI alleles had an additive effect. Using a linear model to predict age of diagnosis as a function of an individual’s total number of autoimmune alleles, the discovery cohort showed a significant 2.6 delayed years to melanoma diagnosis per autoimmune allele (**Fig. 1D**, p = 0.002; CI = [0.935, 4.262]).

To confirm the generality of these findings, we investigated the age of melanoma diagnosis and autoimmune allele presence in an independent validation cohort of 586 individuals diagnosed with cutaneous melanoma, compiled from 4 published dbGaP studies ^92–96^ and the UKBioBank. Despite our best efforts to construct a validation cohort similar to our discovery cohort, we observed notable differences in the distribution of sex and age (**Supplementary Table 1**), especially for the UKBioBank. In contrast to TCGA, females in the validation cohort were significantly associated with an earlier age of melanoma diagnosis independent of AI status (**Supplementary Fig. 5A**; p = 0.0033, median earlier age of diagnosis difference = 3 years). Given not only the existence, but also the direction of observed sex-specific age effects (i.e, females exhibiting an earlier age of diagnosis, rather than males), this potentially represents an intrinsic selection bias within the studies from which our validation cohort was compiled. Despite this, we again observed that having at least one autoimmune-linked MHC-I allele was significantly associated with a later age of diagnosis (**Fig. 1E**, p = 0.0435; median age separation = 1 year). While the direction of this effect was conserved across males with a median age difference of 1 year, we did not observe any significant differences in females (**Supplementary Fig. 5B**; p_male_ = 0.055, p_female_= 0.236).

As most validation samples lacked tumor sequencing data, it was not possible to estimate UV exposure. Thus, our regression analysis was limited to presence of an AI allele and sex. Validation individuals with at least one AI allele had a predicted 2.10 delayed years to melanoma diagnosis relative to those without any of these alleles (p = 0.082; CI = [-0.265, 4.457]). By comparison, the discovery cohort showed a predicted 4.0 delayed years to melanoma diagnosis when only AI status and sex are considered (p = 0.006; CI = [1.154, 6.850]). In the 559 validation cohort individuals with fully-resolved HLA types, we observed a similar trend for total AI allele burden to associate with later age at diagnosis, with a predicted 0.727 year delay to melanoma diagnosis per autoimmune allele, though these results did not reach statistical significance (**Fig. 1F**, p = 0.328; CI = [-0.731, 2.184]).

Finally, we attempted to identify other HLA alleles associated with age at diagnosis by comparing individuals with a particular allele to all individuals without that allele. While none of the associations were significant after multiple hypothesis testing correction, we did note that HLA-B*27:02, showed a marked effect across cohorts with an earlier median age of diagnosis of 10 years in the SKCM-TCGA (15 individuals; p = 0.0496; p_adj_ = 0.483) and 11 years in the validation cohort (3 individuals; p = 0.0332; p_adj_ = 0.388) respectively (**Supplementary Fig. 6**). However, given the limited cohort sizes with sequencing data, coupled with the highly polymorphic nature of the HLA, this analysis is underpowered and may warrant further investigation in larger cohorts.

### MHC-I Autoimmune Alleles Modify Melanoma Risk

Polygenic risk scores (PRS) use information about genetic risk factors to predict individual disease risk. We evaluated the utility of incorporating MHC-I AI allele carrier status into a risk scoring framework in the context of the PRS developed by Gu *et al.*^97^ (Methods: *PRS Implementation)*. This PRS comprises 204 SNPs, of which 16 are found on chromosome 6, though none fall within the HLA class-I region (**Fig. 2A**). Although the HLA class-I genes are not among the genes associated with each SNP as reported by Gu *et al.*,^97^ we assessed whether HLA-proximal PRS SNPs (i.e., those SNPs within 3MB of the HLA-coding region) associated with MHC-I AI allele carrier status across our discovery cohort, but observed no significant relationship (p = 0.24).

**Figure 2:**
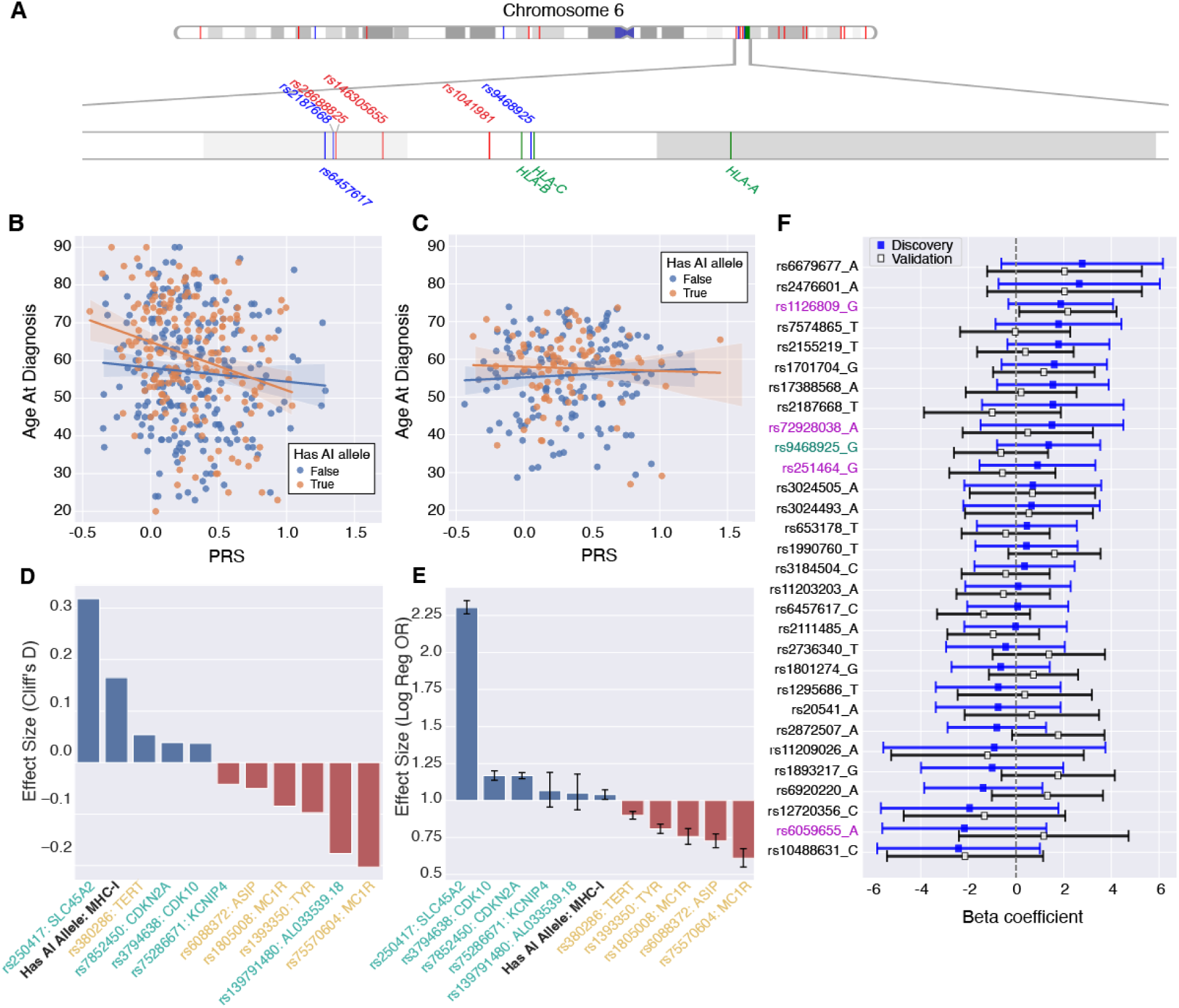
A) A PhenoGram^99^ plot of chromosome 6 shows melanoma PRS SNPs (red) and autoimmune-associated SNPs (blue) fall outside of the HLA coding region (green). The distance from the closest PRS SNP (rs1041981) to the class-I coding region (HLA-B) is 215,819 bp. The closest AI SNP (rs9468925) falls in between HLA-C/HLA-B. B) *TCGA:* Age of diagnosis as a function of PRS and autoimmune allele presence. Individuals with autoimmune MHC-I alleles show a steeper decrease in age of diagnosis as PRS increases (p = 0.055, Beta = -9.124, CI = [-18.432, 0.183]). C) *Validation:* Age of diagnosis as a function of PRS and autoimmune allele presence. Individuals with autoimmune MHC-I alleles again show a steeper decrease in age of diagnosis as PRS increases, but interaction effects were insignificant (p = 0.499, Beta = -2.878, CI = [-11.254, 5.498]). D) *TCGA:* Cliff’s D for having at least one MHC-I linked autoimmune allele, the 5 strongest PRS melanoma predisposing SNPs mapped to their nearest gene (yellow), and the 5 strongest PRS melanoma protective SNPs mapped to their nearest gene (turquoise). E) *MVP:* Inverse odds ratios (1/OR) for MHC-I AI allele presence, the 5 strongest PRS melanoma predisposing SNPs mapped to their nearest gene (yellow), and the 5 strongest PRS melanoma protective SNPs mapped to their nearest gene (turquoise). F) *Autoimmune SNP effect on age of diagnosis*: Autoimmune SNPs showed varying associations with age of diagnosis ranging from strongly protective (high Beta) to strongly predisposing (low Beta). Overall though these effects were highly variable and did not reach statistical significance after multiple-hypothesis correction. Joint vitiligo-melanoma associated SNPs are marked in red, the lone HLA-C/HLA-B psoriasis and vitiligo SNP is marked in green, and the remaining 25 broad autoimmune SNPs are marked in black. Error bars correspond to +/- 2 standard-deviations.

As the PRS was developed to stratify cases from controls, we ensured that it could be generalized to capture age-specific effects in a case-only cohort through regression analysis. We observed that higher PRS associated significantly with earlier age of diagnosis in the TCGA (*Discovery*: p = 0.002, Beta = -7.499, CI = [-12.141, -2.857]), but not in the validation cohort (*Validation*: p = 0.710, Beta = 0.7551, CI = [-3.237, 4.747]). Given the disparity in PRS generalization, we evaluated the PRS age stratification in melanoma cases from the Melanostrum Consortium (N = 3001), the original PRS validation set.^97^ Here we observed that higher risk scores were associated with an earlier age of diagnosis (p = 0.055, Beta = -1.780, CI = [-3.598, 0.038]). We compared the minor allele frequency (MAF) of risk SNPs across these three datasets and observed strong correlations. However, we did observe certain SNPs with a MAF difference ≥ 0.1 across datasets. Four SNPs exhibited this difference between our discovery set and Melanostrum (rs1464510, rs187989493, rs7041168, rs7164220). Between our validation set and Melanostrum, seven SNPs exhibited this MAF gap (rs187989493, rs1393350, rs7164220, rs13338146, rs12919293, rs75570604, rs2092180) including the maximum PRS effect SNP (rs75570604). Finally, between our discovery and validation sets one SNP exhibited a MAF difference ≥ 0.1 (rs1464510) (**Supplementary Fig. 7A-C**). These discrepancies may explain the varying performance of PRS in age stratification across cohorts.

We next evaluated the relationship between PRS and age at diagnosis in AI allele carriers versus non-carriers. As expected, PRS score distributions did not differ between those with and without MHC-I AI alleles in either cohort (**Supplementary Fig. 8A-D**; p_disc_ = 0.999, p_val_= 0.901). Interestingly we observed that increases in PRS showed a greater negative effect on age at diagnosis in those with an autoimmune allele relative to those without one in the discovery cohort (**Fig. 2B**). Interaction effects from a linear model between PRS and AI MHC-I allele genotype trended negatively (p = 0.055, Beta = -9.124, CI = [-18.432, 0.183]). This could suggest that interactions between known risk SNPs and MHC-I genotypes play a role in melanoma predisposition, or that known high risk variants can overwhelm the protective effect of carrying an MHC-I AI allele. We did not observe any significant interaction effects between PRS and AI MHC-I allele genotype in the validation cohort (**Fig. 2C**).

We also sought to assess whether these findings generalized to melanoma incidence. In a large case-control cohort composed of individuals from the Million Veteran Program^98^ (MVP; N = 187,292; *Methods: Datasets*), AI MHC-I carriers were less likely to have a melanoma diagnosis (OR = 0.962, p = 0.024), as would be expected if MHC-I AI alleles conferred a protective effect against melanomagenesis. To quantify the magnitude of AI MHC-I allele carrier status relative to individual PRS SNPs, we compared effect sizes in both the discovery cohort and the MVP. PRS stratification in the MVP yielded an AUC of 0.61, in line with, but less than the reported 0.644 AUC in the Melanostrum Consortium ^97^ (Methods: *PRS Implementation*). Compared to the five largest PRS weight SNPs in both directions (i.e., melanoma predisposing and protective), MHC-I AI allele status exhibited the fourth largest effect size by magnitude and second largest positive effect size in the discovery cohort (**Fig. 2D**, Cliff’s D = 0.165). In the MVP, the effect of having an MHC-I linked AI allele was more modest, with the smallest odds ratio by magnitude relative to the 10 largest weight PRS SNPs (**Fig. 2E**, OR = 0.962, CI = [0.930,0.995).

We next evaluated whether these trends extended to non-MHC AI risk SNPs. In total we examined 30 AI SNPs, including four with established vitiligo-melanoma associations either as the joint lead risk SNP for both conditions (rs1126809, rs6059655) or in strong linkage disequilibrium with known cutaneous melanoma risk SNPs (rs72928038, rs251464),^30^ and one (rs9468925) that is associated with both psoriasis and vitiligo and falls in between HLA-C/HLA-B.^60^ The remaining 25 AI SNPs are broadly associated with autoimmunity (i.e., associated with at least three autoimmune conditions, and at least one of which surpassed a GWAS significance of p = 10^-7^) and were previously investigated in the context of immune-checkpoint inhibitor success in melanoma by Chat *et al*.^38^ While coefficients for the relationship between AI SNP genotype and age of melanoma diagnosis ranged from strongly protective (e.g., rs6679677; Beta_Disc_ = 2.775; Beta_Val_ = 2.036) to strongly predisposing (e.g., rs10488631; Beta_Disc_ = -2.407; Beta_Val_ = -2.137) across cohorts, overall these effects exhibited large variability and were not significant after multiple-hypothesis testing (**Fig. 2F**). In contrast to MHC-I AI alleles, including non-MHC-I AI SNPs as covariates with PRS did not improve prediction of age at diagnosis.

While cancer cohorts document age at diagnosis, the ideal value to consider for a risk analysis is age at onset, as this would suggest optimal screening times for early detection. To estimate the potential “window of opportunity” for earlier melanoma detection, we estimated the expected time between the initial transformed malignant cell (in a surviving malignant clone that escapes extinction) and clinical detection using a multistage carcinogenesis model for melanoma (Methods: *Multistage Carcinogenesis Model for Melanoma*). Briefly, we developed a cell-based stochastic branching process model for the development of independent premalignant clones (such as nevi) that can arise and clonally expand in normal skin epithelium. Each cell in these clones has the propensity to transform to a malignant cell with a certain probability, the malignant clone population can expand in size or go extinct through a stochastic birth-death process, and clinical detection may occur with a size-based detection probability. Mathematically, the expectation of the lag-time variable, or the time between the founder cell of a persistent malignant clone and clinical detection, can be interpreted as the average “age” or sojourn time of the detected tumor.^100^

We analyzed Surveillance, Epidemiology, and End Results (SEER9) melanoma age- and cohort-specific incidence data from 1975-2018.^101^ The hazard function from a “two-stage” model corresponded to the best fit to SEER incidence for both men and women (**Supplementary Fig. 9**). Adding additional stages to the model (i.e., more than 2 rate-limiting events or “hits” such as driver mutations required before malignant transformation) did not improve the fits. With estimated model parameters, we found that the expected tumor sojourn time in males was 8.35 years (Markov chain Monte Carlo [MCMC] 95% CI: 6.61 - 9.73) and similarly in females was 9.64 years (95% CI: 8.48 - 10.66). Previous studies have estimated melanoma doubling times that can be used to then calculate the corresponding tumor sojourn time. With a mean doubling time of 144 days,^102^ the mean growth rate in an exponentially growing tumor is approximately 1.76 per year. Assuming malignant tumors are detected on average at 10^8^ or 10^9^ cells in size, this implies a melanoma sojourn time of 10.5 years and 11.8 years, respectively. This is in line with our above estimates, along with those found in a previous modeling study of melanoma doubling times (mean = 3.78 months).^103^ Although estimates may vary based on patient-specific factors, our findings suggest that it takes approximately a decade on average for a melanoma to be detected after it is first initiated in an individual.

Subtracting the ∼10 year sojourn time from age of diagnosis, we further partitioned our discovery cohort into PRS quintiles and stratified by AI-carrier status. AI carrier status exhibited significant later predicted ages of onset for the lowest (p = 0.005) and second-highest risk quintiles (p = 0.034) (**Supplementary Fig. 8E**). Across quintiles we observed the median predicted onset age ranged from 44-55, within the proposed melanoma screening range. In the future customizing melanoma onset estimates from tumor genetic and epigenetic marks has the potential to improve screening approaches based on germline genetic risk factors.

### Investigating Autoimmune Alleles in Melanoma ICPI Response

Immune-checkpoint inhibitors (ICPI) induce immune activation and stimulate anti-cancer responses through blockade of the inhibitory proteins CTLA-4 (cytotoxic T lymphocyte-associated protein-4), PD-1 (programmed cell death protein-1), and PD-L1 (programmed death-ligand 1). While ICPIs have revolutionized cancer therapeutics, individual responses are highly variable. While long-term durable clinical responses have been documented, many individuals either fail to respond or instead progress and treatment can cause severe immune-related adverse events (irAEs). This has motivated efforts to stratify likely responders before ICPI administration. Melanoma is one of the best ICPI responders across cancer types with estimated response rates of 20% for anti-CTLA4 monotherapy,^104,105^ 30-40% for anti-PD-1 or PD-L1 monotherapy,^106–108^ and 60% for combination therapy.^108,109^ Biomarkers including tumor mutation burden and PD-L1 positivity have been found to associate with response, however, models based on these markers still have significant false positive and negative rates. Given the importance of neoepitope presentation in ICPI responses, it is likely that MHC-genotype plays a role as well.^49–51^

Interestingly, vitiligo as an irAE to ICPI administration is associated with improved prognosis and tumor regression.^63–66^ If ICPIs can induce remission through broad melanocytic destruction mediated by CD8+ T-cells, then the presence of MHC-I AI predisposing alleles may have a role in pre-treatment stratification. Therefore, we investigated whether any relationship existed between our set of MHC-I AI alleles and clinical ICPI response in melanoma using a subset of our validation cohort (N_anti-CTLA4_ = 103, N_anti-PD-1_ = 35). Response was characterized in accordance with the methodology of the original studies. Specifically, those with (ir)RECIST criteria of either complete response, partial response, or stable disease with an overall survival exceeding one year were labeled as responders (N_responders_ = 45). Overall, we did not observe any significant associations between MHC-I AI allele carrier status with ICPI response (OR = 0.69, p = 0.36) or with HLA-A*02:01 carrier status either in isolation (OR = 1.06, p = 1.0) or in the context of broad MHC-I AI allele carrier status (OR = 1.15, p = 0.85). Granular ICPI separation to a treatment-specific level of anti-CTLA4 (OR = 0.59, p = 0.36) or anti-PD1 (OR = 0.97, p = 1.0) similarly did not show significant associations.

### Investigating Mechanisms of MHC-I Autoimmune Allele Protection

One possible explanation for a protective effect of selected HLA alleles would be a stronger affinity for neoantigens or tumor associated antigens that cannot easily be suppressed by melanomas. In psoriasis, melanocyte antigens such as ADAMTS-like protein 5 presented by HLA-C*06:02 (one of the seven AI alleles) can induce a targeted CD8+ T-cell response against melanocytes.^59^ Similarly, in vitiligo, CD8+ T-cells target antigens from melanosomal proteins such as PMEL, MART1/MLANA, TYR, TRP-1, and TRP-2.^76,110,111^ Interestingly melanoma-specific CD8+ T-cells appear to recognize peptides derived from these conserved melanocytic antigens more often than melanoma-specific antigens.^76,112–114^ Given this, a protective effect in cancer could indicate that AI alleles mediate more effective immune surveillance against conserved cancer antigens (*i.e.*, self-antigens overexpressed in tumors) or even against somatic mutations that promote tumor development.

We reasoned that a protective effect manifesting as delayed age of diagnosis would require more effective immune surveillance against early driver mutations or conserved melanocyte-specific antigens expressed by melanomas. For neoantigens, we identified a set of 215 mutations that exhibited joint high DNA and RNA variant allelic fraction coupled with either recurrence or driver likelihood as predicted by the CHASM algorithm ^115,116^ (Methods: *Identifying Driver Mutations*). This included well-known driver mutations in BRAF, NRAS and CDKN2A (**Fig. 3A-B**), with BRAF V600E being the most frequent mutation across the discovery cohort at 164 unique occurrences (**Fig. 3A**). For conserved antigens, we included peptides derived from antigens associated with melanocytes ^76,110^ and melanoma, including the melanoma antigen gene (MAGE) family.^117–123^ We also expanded our list of conserved antigens to include genes constitutively expressed in melanocytes (Methods: *Identification and Differential Expression of Conserved Antigens*). In total, we considered 91 genes as sources of conserved antigens (**Supplementary Table 3**). Transitioning from identification to investigation, we evaluated whether MHC-I AI alleles could better expose driver neoantigens and conserved antigens for immune surveillance than 2,908 other common MHC-I alleles based on NetMHCPan-4.1 binding affinity scores for 8-11mer peptides, using 0.5 and 2 percentiles as strong and weak binding cutoffs respectively (Methods: *Predicting Binding Affinities*).^124^

**Figure 3:**
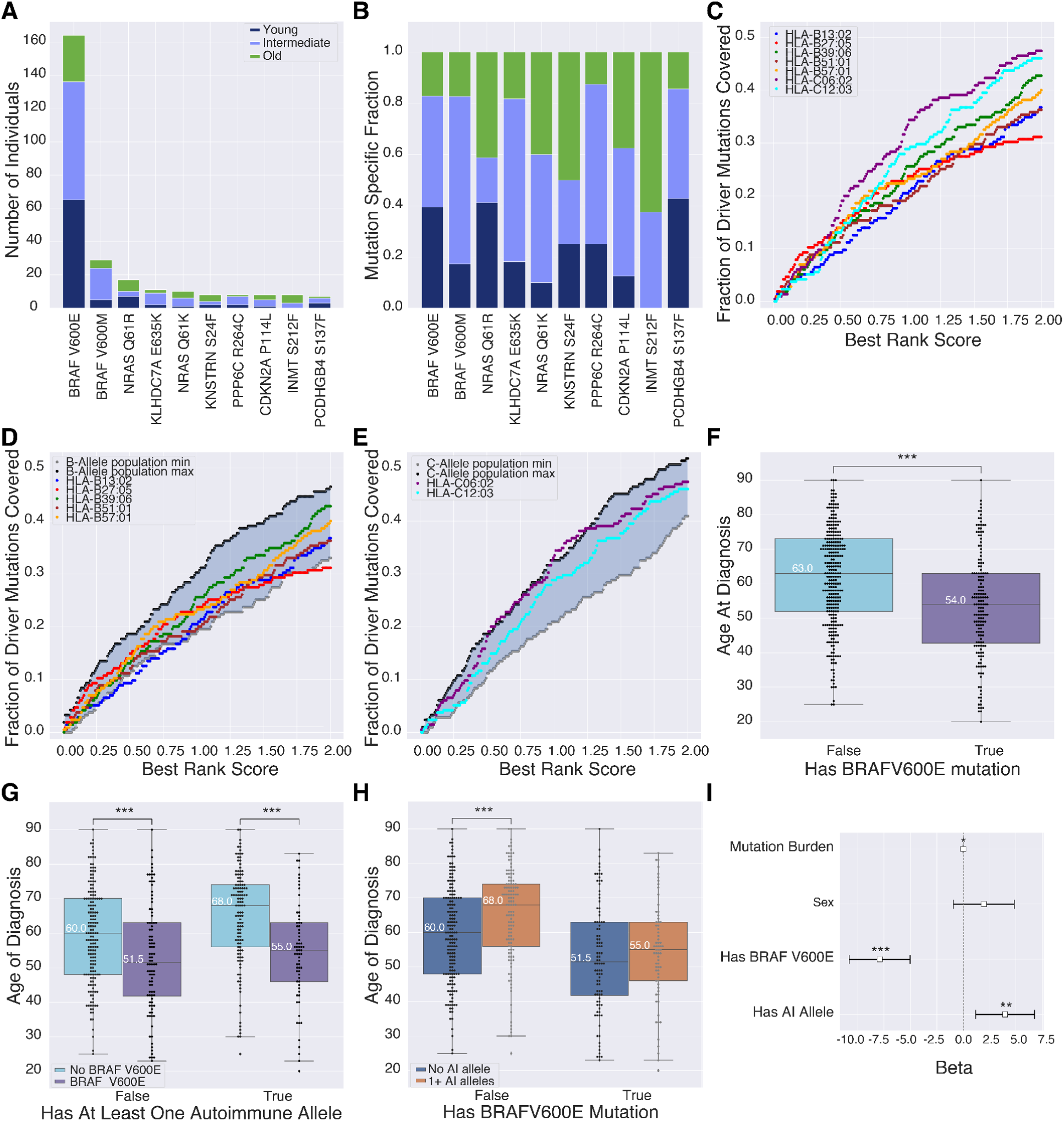
A) Frequency of the 10 most recurrent mutations across the TCGA by age group. Young (< 50) and old (≥ 69) age groups correspond to the bottom and top 30% of individuals by age respectively. The intermediate (50 ≤ x < 69) age group corresponding to the remaining 40% of individuals. B) Relative age group distribution of the 10 most recurrent mutations across the TCGA. C) Fraction of driver mutations presented by autoimmune alleles as a function of BR score. D) Fraction of driver mutations presented by HLA-B autoimmune alleles relative to common (≥ 1% population frequency; 19 alleles) HLA-B alleles. Maximum and minimum population allele coverage corresponds to the maximum and minimum fraction of driver mutations capable of being presented across common alleles at each best rank score respectively. E) Fraction of driver mutations presented by HLA-C autoimmune alleles relative to common (≥ 1% population frequency; 13 alleles) HLA-C alleles. F) Individuals with a BRAF V600E mutation show a significant 9 year earlier age of melanoma diagnosis relative to those without this mutation (p = 7.58x10^-6^). G) BRAF V600E significantly reduces melanoma age of diagnosis across discovery cohort individuals independent of autoimmune allele presence with median earlier ages of diagnosis of 8.5 years in those without an AI allele (p = 7.39x10^-4^) and 13 years in those with an AI allele (p = 8.12x10^-7^). H) Discovery cohort individuals with an AI allele show a significant median later age of diagnosis of 8 years in the absence of a BRAF V600E mutation (p = 2.98x10^-4^). However BRAF V600E mutation presence appears to counter the AI allele protective effect with a loss of significance between those with and without AI alleles (p = 0.232) I) BRAF V600E mutation presence is significantly associated with 7.91 earlier years to melanoma diagnosis (p_BRAFV600E_ = 7.12x10^-8^), while having at least one MHC-I linked autoimmune allele is significantly associated with 3.97 delayed years to melanoma diagnosis (p_autoimmune_ = 4.59x10^-3^) after controlling for sex and mutation burden. Mutation burden also remained significant, but had a minimal contribution to age of diagnosis with 0.003 delayed years until melanoma diagnosis (p_mutation_ = 0.014)

There was variability in the potential of MHC-I AI alleles to present peptides containing driver neoantigens in general, with HLA-C*06:02 and HLA-C*12:03 presenting the largest proportion of mutations across all affinities, and HLA-B*27:05 presenting the largest proportion at stronger affinities (**Fig. 3C**). As compared to common alleles (Methods: *HLA Population Allele Representations)*, MHC-I AI alleles did not show particularly better binding affinity for neoantigens for either HLA-B or HLA-C alleles (**Fig. 3D-E**), though HLA-C*06:02 had coverage close to and briefly exceeding the common allele maximum (**Fig. 3E**). In general, maximum and minimum population B-alleles exhibited greater coverage than their corresponding C-alleles at lower rank scores, but C-alleles outperformed B-alleles as the rank score exceeded 0.5 (**Supplementary Fig. 10A**).

There were also differences in which mutations generated neopeptides with the best specificity. While common and AI alleles on average exhibited specific mutation binding preferences (**Supplementary Fig. 11**) among the 215 drivers, there was no single mutation that was more effectively presented by all MHC-I AI alleles versus common alleles. One mutation, BRAF V600E, showed significant age differences with mutation carriers being diagnosed with melanoma on average 9 years earlier than those without (**Fig. 3F**; p = 7.58x10^-6^). However, rather than observing a correlation between lack of AI allele presence and having a BRAF V600E mutation, BRAF V600E status served as an indiscriminate melanoma catalyst, significantly shifting age of diagnosis earlier regardless of AI allele status (**Fig. 3G**). Moreover, it appeared to counter the AI allele protective effect with a reduced age gap between those with and without AI alleles in BRAF V600E tumors (**Fig. 3H**). Regression analysis showed similar results with V600E mutation presence having a larger effect size than AI carrier status by almost 4 years. (**Fig. 3I**).

Across AI alleles, only three were predicted to present BRAF V600E. HLA-B*27:05 was the only allele with a predicted affinity below the strong binding cutoff (best rank score = 0.22). HLA-B*27:05-restricted cytotoxic T-cell responses have been observed against V600E.^125^ HLA-B*39:06 and HLA-B*57:01 had scores of 1.78 and 0.61 respectively, showing potential for weaker V600E binding. We found no association with occurrence (OR = 1.04, p = 0.84) or expression of BRAF V600E (p = 0.34) in individuals carrying AI alleles in general, or those carrying one or more of these three AI alleles specifically. An association might have been expected if the mutant allele was subject to strong counter selection by immune surveillance. Notably, BRAF V600E has been suggested to avoid immune surveillance by accelerating internalization of cell surface MHC-I.^126^

We next compared affinity of MHC-I AI alleles for conserved antigens to common alleles. Here, we considered differences in NetMHCPan-4.1 affinities both regionally and on average for a set of 91 proteins, including known melanoma cancer antigens^76,110, 117–123^ and an additional 52 genes that were both stably and specifically expressed in melanocytes and were expressed in melanomas (**Supplementary Table 3).** These 52 included four well-known melanocyte genes, PMEL, MLANA, TYRP1, and TYR, and the rest were enriched for the folate metabolism pathway which is important for DNA repair in melanocytes.^127^ Evaluating expression in melanomas relative to melanocytes we observed that two MAGE genes, MAGEA10 and MAGEE1, were significantly upregulated and that several canonical melanocyte and stably expressed genes were downregulated (**Fig. 4A**). This is consistent with reports that MAGE genes are specific to reproductive tissues ^128^ and tumors ^117–123^ and canonical melanocyte genes such as TYRP1 are minimally expressed, if not undetectable, in melanoma.^113,129,130^

**Figure 4:**
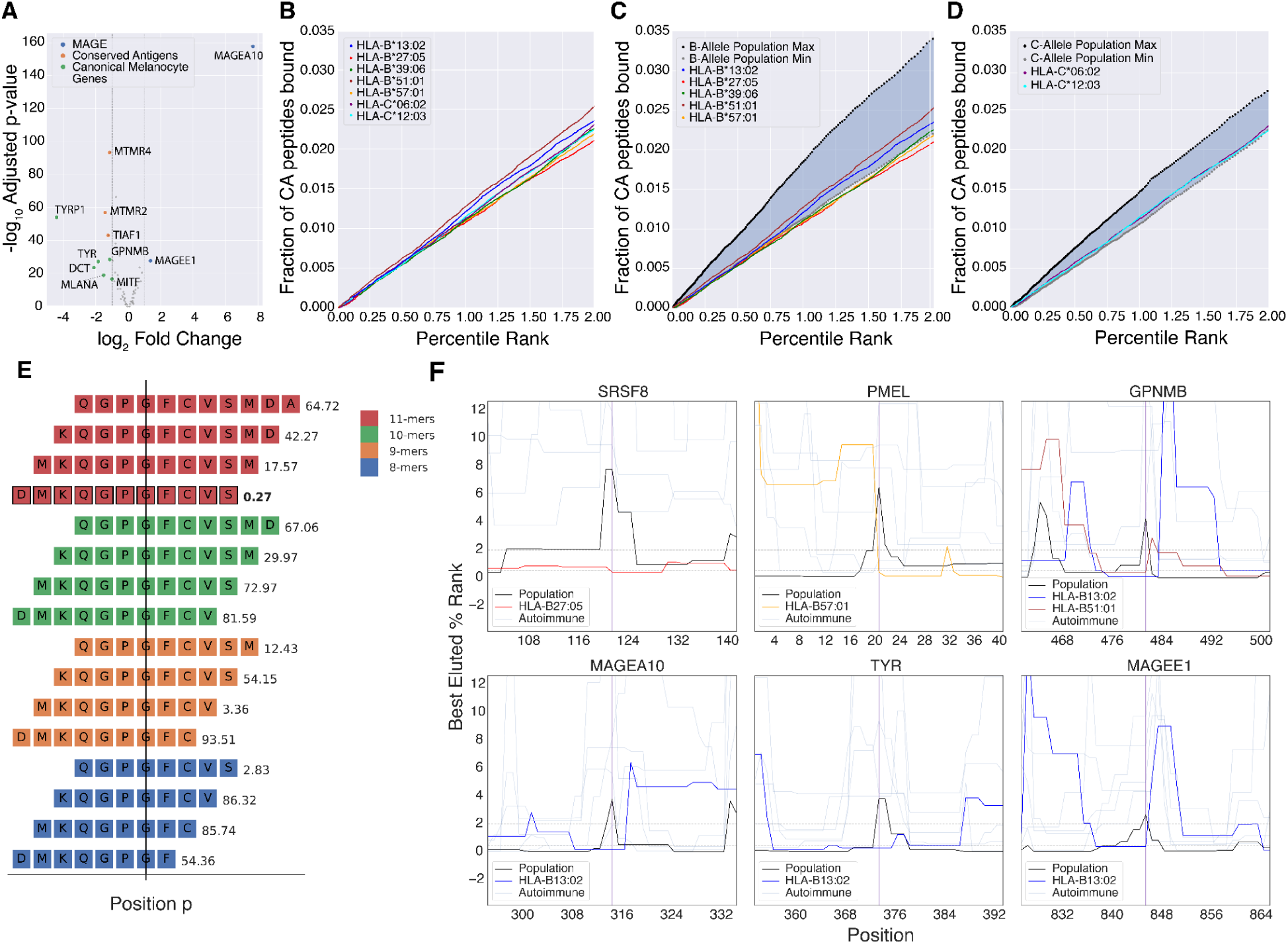
A) Differential expression of conserved antigens between normal melanocytes and melanoma. Labeled genes have an adjusted p-value > 0.5 and an absolute fold change > 2. B) Fraction of peptides from conserved antigens (CA) presented by each autoimmune allele. C) Fraction of peptides from conserved antigens (CA) presented by HLA-B autoimmune alleles relative to common HLA-B alleles (≥ 1% population frequency; 19 alleles). Maximum and minimum population allele representations correspond to the maximum and minimum fraction of conserved antigen peptides bound across common alleles at each rank score respectively. D) Fraction of peptides from conserved antigens (CA) presented by HLA-C autoimmune alleles relative to common HLA-C alleles (≥ 1% population frequency; 13 alleles). E) Position-wise best percentile rank scores were computed from predictions for 8-11mers by taking the best percentile rank of all overlapping peptides at any given position. F) Greatest position-wise differences in best percentile rank between HLA-B autoimmune and common alleles amongst the top 2 differences (*SRSF8* and the canonical melanocyte gene *PMEL*) and several DE conserved antigens (*GPNMB, MAGEA10, TYR, MAGEE1*). Common alleles are shown in orange and autoimmune alleles are shown in transparent blue unless they are predicted to elute at better percentile ranks than common alleles. Vertical purple lines demarcate positions where autoimmune alleles are predicted to elute at better percentile ranks than common alleles; plots show up to +/- 20 amino acids of the demarcated positions.

To evaluate broad allele-specific differences in conserved antigen-derived peptide repertoires, we compared the fraction of 8-11mers from each conserved antigen predicted to bind at a given percentile rank (Methods: *Predicting Binding Affinities*). Across AI alleles we observed minimal variation with HLA-B*51:01 being the most promiscuous AI allele and HLA-B*13:02 being the second most promiscuous across the binding range (**Fig. 4B**). However, in general, AI alleles exhibited a narrower conserved antigen repertoire relative to common alleles (**Fig. 4C-D**). HLA-B though generally presented a broader range of peptides across conserved antigens than HLA-C (**Supplementary Fig. 10B**). Considering individual conserved genes, AI alleles were generally predicted to bind a smaller fraction of derived peptides than common alleles. Comparison of the distributions of the top 10 differences by mean at a gene-specific level showed that almost all MHC-I AI alleles present conserved antigens no better than common alleles (**Supplementary Fig. 12**), with the notable exception of HLA-B*27:05’s affinity for SRSF8 and TOMM6 (**Supplementary Fig. 12A-B**).

Since MHC-presented peptides are not uniformly sampled across the entire parent protein^131^ we revisited our analysis with a focus on regional differences by calculating position-wise best percentile ranks across all 8-11mers overlapping a position in each gene (**Fig. 4E**). We did not observe any region of a conserved antigen that was consistently more effectively presented by MHC-I AI alleles relative to common alleles, however there were short regions where specific MHC-I AI alleles had stronger affinity than common alleles (**Fig. 4F**). In addition to binding SRSF8 more generally, HLA-B*27:05 also exhibited the greatest position-wise advantage to SRSF8 over common alleles (**Fig. 4F**). This is in spite of having the smallest conserved antigen repertoire overall (**Fig. 4B-C**). Several proteins had regions with affinity differences between HLA-B AI and common alleles, specifically the canonical melanocyte genes *PMEL, GPNMB*, *TYR*, and *MITF* and the MAGE genes *MAGEA10 and MAGEE1* (**Fig. 4F, Supplementary Fig. 13**). However, regional affinity advantages of AI alleles over common alleles tended to be allele, gene, and position specific with little overlap across AI alleles. Overall, these analyses suggest that the protective effect of MHC-I AI alleles does not derive from a shared specificity for a conserved antigen or neoantigen, but cannot rule out that differences in specificity contribute to the effect.

## Discussion

Immunosurveillance has been implicated in melanomagenesis prevention,^16,17,31^ yet HLA contributions to melanoma risk have largely remained uncharacterized. Here we investigated whether predisposing MHC-I alleles for CD8+ T-cell driven skin-associated autoimmune disorders (vitiligo and psoriasis) could protect against melanoma. Our findings support this hypothesis, with AI-allele carriers exhibiting a significant later age of melanoma diagnosis in both the TCGA and an independent validation cohort. Moreover, AI-allele specific protection appears not only to be uncaptured by current melanoma PRS,^97^ but also can augment PRS performance in relative risk stratification. While at least 6 vitiligo risk SNPs are protective from melanoma,^28,29^ our results show that autoimmune risk effects can be extended to MHC-I as well, and suggest a broader space of joint autoimmune predisposition and melanoma protection.

We further investigated potential mechanisms linking MHC-I AI alleles to delayed age at diagnosis. Immune activity against melanoma has been linked to high mutation burden due to UV exposure,^21,22^ suggesting neoantigens could provide a potential substrate. Focusing on mutations that drive melanomagenesis, we did not see obvious differences in the affinity for MHC-I AI alleles for neopeptides relative to other alleles. Interestingly, lack of AI allele association with improved response to immunotherapy would seem to support that neoantigens are not the mechanism by which AI alleles boost immune surveillance in melanoma. While AI alleles did not seem to interact with specific mutations, we did note that BRAF V600E was associated with an earlier age at diagnosis independent of AI allele carrier status. Potential of autoimmune alleles to present BRAFV600E did not appear to impact the incidence of the mutation, which is consistent with reports that this mutation associates with impaired immune surveillance.^126^

As CD8+ T-cells target healthy melanocytes through conserved antigens in both psoriasis and vitiligo, responses against conserved antigens could also provide an explanation. We observed positions with greater presentability by AI alleles in genes spanning all subcategories of conserved antigens including canonical melanocyte genes, melanoma antigen genes, and genes stably expressed in melanocytes. In particular, AI alleles had greater presentability of amino acids in PMEL and TYR (**Fig. 4F**), which are melanocyte-specific antigens recognized by CD8+ T-cells in melanoma-associated vitiligo.^76,110^ However, we did not observe position-wise advantages for AI alleles in other CD8+ T-cell targets reported in melanoma-associated vitiligo, such as MLANA or TYRP2. MHC-I AI alleles also uniquely target positions in the melanoma antigen genes MAGEA10 and MAGEE1 which are upregulated in melanoma relative to melanocytes (**Fig. 4A,F**). Finally, we observed positions uniquely targeted by AI alleles within melanocyte stably expressed genes. Notably, SRSF8 exhibited the position most favored by AI alleles and is overall substantially better targeted by the AI allele, HLA-B*27:05 (**Supplementary Fig. 12**). This suggests that melanocyte-specific conserved antigens other than those already identified in melanoma-associated vitiligo may also contribute to the protective effect of AI alleles. Additionally, while canonical melanocyte genes have been known to be recognized by CD8+ T cells, they are typically downregulated in melanoma (**Fig. 4A**) or otherwise inconsistently expressed.^132,133^ In contrast, the novel conserved antigens described in this study exhibit stable expression across melanocytes and melanoma and might serve as more consistent immune targets.

Altogether, these findings support that AI alleles’ unique immunopeptidomes could contribute to their protective effect against melanoma. However, further investigation is needed. In particular, some studies have suggested that stability of the MHC-I-peptide complex distinguishes AI alleles from other MHC-I alleles,^134–136^ so it is possible that the mechanism is not fully dependent on antigen specificity. We noted that protective effects were confined to 7 specific AI-associated HLA alleles, and were not shared by broader HLA supertype groupings supporting that allele-specific characteristics, whether related to unique antigen specificities, stability or some other characteristic, are likely to account for any protective effect.

In general, AI alleles presented a narrower peptide repertoire compared to common alleles (**Fig. 4C-D**). Notably, HLA-B*27:05 and HLA-B*57:01, both of which covered a smaller fraction of potential conserved peptides than the minimum HLA-B population allele representation, are known for their fastidiousness, or narrow peptide-binding repertoire, as studied in the context of progression from HIV infection to AIDS.^137^ Košmrlj *et al*. also observed that fastidious alleles present peptides not found in common alleles’ repertoires. We similarly observed that the most fastidious AI alleles, HLA-B*27:05 and HLA-B*57:01, exhibited the greatest position-wise advantages over common alleles. Additionally, fastidious class I alleles are expressed on the cell-surface at much higher levels than their promiscuous class I counterparts,^138^ and therefore likely offer more opportunities for both neoepitope and self-antigen presentation from their binding repertoires. Interestingly, we note that outside of autoimmune associations four of our seven AI-alleles (HLA-B*27:05, HLA-B*51:01, HLA-C*06:02 and HLA-B*57:01) are among the strongest HIV-protective alleles.^70,139,140^

Peptide-MHC (pMHC) affinity is driven largely by MHC-I sequence variation where specific polymorphisms shape the binding pockets in the peptide-binding groove. Unsurprisingly, given the skin-specific autoimmune associations for which these alleles were selected, certain AI-alleles have similar binding pocket characteristics. For example, HLA-C*06:02 and HLA-C*12:03 both share a strongly negative E-pocket.^141^ HLA-C*06:02 shares an electronegative B-pocket with HLA-B*27:05 as well.^141,142^ An allele’s affinity-based presentable peptide repertoire though is an idealistic representation that fails to account for antigen processing pathway contributions. Ultimately the space of bound peptides presented on the cell-surface is far narrower. Endoplasmic reticulum aminopeptidases (ERAP) 1 and 2 play an essential role in MHC-I antigen processing. They function to clip peptides to the appropriate size for MHC-I binding,^143,144^ but can also destroy potential ligands through overtrimming.^145–148^ ERAP1 and ERAP2 are prominent risk factors in MHC-I linked autoimmune conditions and both are associated with psoriasis risk.^149–151^ Epistatic effects between ERAP1 and AI-risk alleles have been observed, particularly with HLA-C*06:02 in psoriasis,^149^ and suggests this risk interaction is tied to an increased likelihood of specific autoantigens making it to the cell surface. Taken together, this suggests skin-specific autoimmune predisposing ERAP genes may also confer melanoma protection and is an interesting area for further pursuit.

Vitiligo has been reported as a favorable irAE correlating with melanoma immunotherapy response.^63–66^ However, we did not observe any associations of AI carrier status with ICPI response. Work by Chowell *et al.*^49^ and Cummings *et al.*^152^ showed that in melanoma the B44 supertype associates with extended survival after treatment with ICPIs. While none of our AI alleles fall within this supertype, we surprisingly observed that the B44 supertype trended towards an earlier age of diagnosis in our discovery cohort (**Supplementary Fig. 4**). The B44 supertype has an electropositive B-pocket, with an affinity for negatively charged residues at P2 such as glutamate (E).^153^ Given this strong glutamate affinity, it may be that in the context of ICPI, the B44 supertype is capable of inducing an immune response through the binding and presentation of the highly recurrent BRAF V600E mutation. In contrast, the MHC-I AI alleles associated with later age at diagnosis do not bind negatively charged residues at P2. In fact HLA-B*27:05 and HLA-C*06:02 have electronegative B-pockets with strong affinities for the positively charged arginine at position 2,^141,142^ and effectively serve as B44 antonyms. This suggests peptide repertoire differences between the B44 supertype and the AI allele set, and further suggests the potential for a dichotomy between MHC-I associated melanoma protection and ICPI response, which is associated with somatic mutation presentation and high tumor mutation burden. We note that sample sizes for assessing the role of AI alleles in ICPI response are small, and revisiting this analysis in larger studies in the future may provide further insight.

In conclusion, our study supports that skin-specific autoimmune MHC alleles have a protective effect in melanoma. We also note the potential for MHC-I mediated autoimmunity to interact with cancer development more broadly. Selecting alleles associated with autoimmune disease affecting the tissue type under investigation may yield similar findings across cancer types. For example, some MHC-I alleles are associated with multiple autoimmune conditions, such as HLA-B*27:05 in both psoriasis and ankylosing spondylitis.^69,70,72,73,154,155^ Ankylosing spondylitis is a targeted form of spinal arthritis primarily affecting the entheses and leads to bone erosion and broad vertebral fusion,^156^ and makes for a conceivable autoimmune counterpart to osteosarcomas. Our study did not address MHC-II alleles which are generally expressed more specifically by antigen presenting cells, but have nonetheless been implicated in both skin autoimmune disorders and immunosurveillance. Taken together, tissue-specific MHC AI carrier status may broaden the scope of the autoimmune-cancer risk interplay and remains an interesting area for further exploration.

## Acknowledgements

This work was supported by a grant from the Harry J. Lloyd Charitable Trust (20191857) to H.C., an Emerging Leader Award from The Mark Foundation for Cancer Research (18-022-ELA) to H.C., a RO1 CA220009 grant to M.Z. and H.C., a R01 MH122688-02 grant to W.K.T., and an NIH (National Institutes of Health) National Library of Medicine training grant (T15LM011271) to A.C. The results shown here are in large part based upon data generated by the TCGA Research Network (https://www.cancer.gov/tcga), the UKBB (Project #37671), and the following studies: phs000452.v2.p1.c2, phs000933.v2.p1.c1, phs001550.v2.p1.c1, phs001500.v1.p1.c1, phs000424.v7.p2, GSE78220. This research also used data from the Million Veteran Program (MVP), Office of Research and Development, Veterans Health Administration. MVP data access was provided under the Genisis Core Project. This publication does not represent the views of the Department of Veterans Affairs or the United States Government.

## Disclosure of Interests

The authors declare no competing interests.

## Author Contributions

Original Concept, J.T., H.C.; Project Supervision, H.C.; Project Planning and Design, J.T., H.C.; Data Acquisition, Processing, and Analysis, J.T., D.L., M.P., A.C.; Melanoma Sojourn Time Estimation, M.L., G.E.L., K.C.; Statistical Advising, W.K.T.; UKBB Data Acquisition, R.M.S.; MVP Data Acquisition and Processing, B.G., C.P, F.N.D, K.M., R.H., S.P, P.S.T; Immunological Interpretation Advising, G.P.M. and M.Z.; Preparation of the Manuscript, J.T., D.L., and H.C.

## Data Availability

All data for this project were obtained from public sources. *Discovery Cohort:* Data were obtained from The Cancer Genome Atlas (TCGA) Research Network (http://cancergenome.nih.gov/). Normal exome sequences and clinical data were downloaded from the GDC on June 23-26^th^, 2018 and April 25^th^, respectively, using the gdc-client v1.3.0. Somatic mutations were accessed from the NCI Genomic Data Commons (https://portal.gdc.cancer.gov/) on May 14^th^, 2017. Genotype calls for TCGA were accessed from GDC on April 26, 2019. *Validation and Stably Expressed Gene Cohorts*: dbGaP data from cohorts phs000452.v2.p1.c2, phs001550.v2.p1.c1, phs000933.v2.p1.c1 and phs001500.v1.p1.c1 were obtained using AsperaConnect v3.9.5.172984. SRA toolkit v2.9.2 was used to obtain WXS/WGS data from the Sequence Read Archive (SRA) (including from the following studies: SRP067938, SRP090294). UKBB data was retrieved under project ID 37671. *Million Veteran Program*: Genotype and phenotype information were obtained through application. *Melanostrum Cohort:* Genotype data from Gu *et al.* ^97^ were obtained by direct communication with the authors.

## Code Availability

Code for all analyses can be found at https://github.com/cartercompbio/MelMHC/

## Methods

### Datasets

TCGA skin cutaneous melanoma tumors (SKCM) were used as the discovery set (N = 470). Cases were retained if they had appropriate clinical information for downstream analysis (i.e., age of diagnosis; 11 did not have this information), were microsatellite-stable (3 MSI tumors were removed), and were at least 20 years of age at time of diagnosis (5 were < 20). Individuals below 20 years of age were excluded due to their increased likelihood of harboring rare predisposing risk variants. After filtering there were 451 tumor samples. MHC-I genotypes were called using the exome based methods POLYSOLVER and HLA-HD.^87,88^

An independent validation cohort of melanoma cases with germline WXS/WGS data was built from 5 separate melanoma studies. Two of these studies (*Hugo et al.* ^93^: SRP067938, SRP090294, *Van Allen et al.* ^92^: phs000452.v2.p1.c2) focused on melanoma response to immune-checkpoint inhibitors (ICPI) and were used to evaluate autoimmune associations in the context of ICPI response. Two of these studies (*Melanoma Exome Sequencing:* phs000933.v2.p1.c1 ^95,96^, *The genetic and transcriptomic evolution of melanoma:* phs001550.v2.p1.c1 ^94^), were broader melanoma studies focusing on the genetic basis of sun-exposed melanoma and melanoma evolution. The final study consisted of individuals from the UKBB with WXS/WGS with ICD10 codes: C433, C434, C436, and C437. Validation cohort individuals were also filtered by age to exclude individuals under 20 years old. MHC-I genotypes for the validation cohort were called using HLA-HD.^88^ Discovery and validation cohorts are described in **Supplementary Table 1**.

Cases from the Melanostrum Consortium (N = 3001) were used to evaluate PRS generalizability from absolute risk (i.e., cases vs. controls) to relative risk (i.e., age-specific effects). Genotype was available at 204 risk SNPs used in the development of the PRS by Gu *et al*.^97^ Cases from this cohort were filtered to include individuals ≥ 20 years in age.

The Million Veteran Program (MVP; N = 187,292) was used to assess the generalizability of our findings to melanoma incidence. Both controls (N = 171,878) and cases (N = 15,414) were constructed from individuals both ≥ 20 years of age at the time of last follow-up and of European descent as determined by HARE (Harmonized Ancestry and Race/Ethnicity).^157^ Controls excluded individuals with non-melanoma cancer diagnoses as determined by PheCode.^158^ MHC-I alleles for the MVP were called using HIBAG^159^ with the multi-ethnic IKMB and Axiom UK Biobank array models.^160^ MVP 1.0 Axiom array design and genotype QC details are described elsewhere.^161^

To identify putative conserved antigens we leveraged RNA-seq from a cohort of healthy melanocytes (dbGaP: phs001500.v1.p1) derived from newborn foreskins (N=106) and version 7 of the Genotype-Tissue Expression (GTEx) project (dbGaP: phs000424.v7.p2).

### Identifying Driver Mutations

We downloaded WES-based mutation calls from the TCGA GDC portal from four different mutation callers: VarScan,^162^ MuSE,^163^ MuTect,^164^ and SomaticSniper.^165^ RNA Variant Allelic Fraction (VAF) was obtained with bam-readcount. From these mutation calls we focused on single nucleotide variants, which dominate the landscape of melanoma and account for the majority of driver events.^166–168^ A mutation was considered to be a potential driver if it: 1) altered protein sequence, 2) was found in both the DNA and RNA in at least one individual, and 3) had a median DNA and RNA variant allele fraction (VAF) percentile less than or equal to 40%. DNA mutations were only considered at a patient-specific level if they were called by at least two of the mutation callers mentioned above. In total 51,062 mutations satisfied these criteria.

We further filtered the list of putative drivers based on recurrence. Specifically, if a specific mutation was detected in 4 or more different tumors we categorized it as a likely driver. In total 109 mutations satisfied this criterion (0.215% of the 51,062 candidate mutations). For those mutations that failed to reach this recurrence, we calculated mutation-specific contributions to melanoma pathogenicity using scores from a melanoma-specific CHASM classifier.^115,116^ Mutations with a CHASM score greater than or equal to 0.9 were deemed to be likely melanoma drivers; 106 mutations satisfied this criterion (0.206% of the 51,062 candidate mutations). Combining these recurrent and predicted driver singleton mutations yielded a final set of 215 melanoma drivers.

### Identification and Differential Expression of Conserved Antigens

GTEx V7 contains 11,688 RNA-seq samples from 714 donors across 53 tissue types and was aligned with STAR v2.4.2a to GENCODE v19 and quantified with RNA-SeQC v1.1.8. RNA-seq reads from healthy melanocytes were aligned with STAR v2.5.0b to GENCODE v19 and quantified with RSEM v1.2.31. After quantification, both cohorts were filtered such that genes with ≤ 0.5 RSEM or with counts < 6 in > 93% of samples were removed. Additionally, ribosomal RNA, Y chromosomal, and histone genes were removed. Ribosomal and histone mRNA are not polyadenylated. Notably, the melanocyte cohort is exclusively male, while GTEx is not, which would potentially lead to Y chromosomal genes being falsely identified as stably expressed downstream.

Across healthy melanocytes and each tissue type in GTEX, genes were scored as stably expressed genes (SEGs) using the output from the scoring method described in scMerge.^169^ Briefly, the method first fits the expression of each gene from each tissue sample to a Gamma-Gaussian mixture. For the expression of a gene *x_i_*, the Gamma component corresponds to samples with low expression and the Gaussian component corresponds to samples with high expression. This mixture has the joint density function

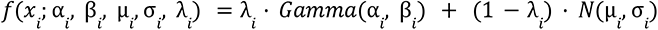

where *α_i_* and *β_i_* are the shape and rate parameters of the gamma component, *μ_i_* and *σ_i_* are the mean and standard deviation of the Gaussian component, and the mixing proportion *λ*_i_ is bounded by [0,1]. The SEG scoring method also takes into account the proportion of zeros in each gene *ω*_i_ for each tissue. Each gene is then scored by the percentile ranks of its mixing proportion *λ_i_*, coefficient of variation (CV) σ_i_/μ_i_, and proportion of zeros ω_i_ uch that the average percentile rank across all three metrics is minimal. The highest scoring genes have lower mixing proportions, CVs, and proportion of zeros.

Using the aforementioned method, each gene from every tissue was fit to a Gamma-Gaussian distribution using scMerge v1.6.0 and given a score from 0 to 1 for stable expression. Then, the set of genes across all GTEx tissues that had scores > 0.69 was removed from the set of genes in melanocytes that had scores > 0.69. The threshold of 0.69 was chosen based on the observation that the scores of canonical melanocyte genes were, with the exception of DCT/TYRP2, all ∼0.7 and by visual inspection of the distributions of scores across tissues (**Supplementary Fig. 14**). This process yielded 4 canonical melanocyte genes (PMEL, MLANA, TYRP1, TYR) and 52 additional protein coding genes for downstream analysis. The 52 non-canonical melanocyte genes were also passed to PANTHER ^170^ for Reactome pathway ^171^ overrepresentation analysis using Fisher’s Exact test, which identified several genes as members of the folate metabolism pathway (FDR 0.0288).

To determine differential expression of conserved antigens in melanoma relative to melanocytes, we applied the same expression quantification pipeline and gene filtering steps to both healthy melanocytes and SKCM. Specifically, we quantified HLA-allele specific expression using HLApers v1.0 ^172^ and the Kallisto v0.44.0 ^173^ pipeline for HLApers. Reads were aligned to GENCODE v30 ^174^ and IMGT HLA v3.41.0.^175^ After quantification, both cohorts were filtered in the same way as the conserved antigen identification pipeline described above. Additionally, 11 samples with missing age of diagnosis were removed from the SKCM cohort. We then performed a differential expression analysis with DESeq2 v1.30.1 ^176^ conditioned on ancestry. However, several potentially relevant covariates were also incompatible across the two cohorts. Namely, age, sex, tumor type (primary or metastatic), tumor purity,^177^ melanocytic plasticity score,^178^ and TIDE score.^179^ Due to these discrepant covariates between the cohorts, we also checked that no covariates were associated with significant DE for any conserved antigens. The only gene subject to DE was MAGEA10 with a -1.3 +/- 0.4 log fold change (LFC) in primary vs. metastatic melanoma. This is substantially less than the +8 LFC of MAGEA10 observed in melanocytes vs. melanoma (**Fig. 4A**).

### Predicting Binding Affinities

MHC-I allele binding affinities were computed across the available 2,915 unique MHC-I alleles for both driver mutations and conserved antigens. Since driver mutations altered protein sequence, we evaluated MHC-I alleles’ ability to present neoepitopes by generating all unique 8-11mers found in a mutation relative to the wild-type (corresponding to the set of novel peptides a MHC-I allele can present to the immune system). To circumvent cross-allele and cross-peptide variabilities that are inherent in predicted IC50 comparisons, we used percentile ranks relative to a random set of peptides provided by NetMHCpan-4.1^180^ to approximate binding affinity for every MHC-I allele peptide pair. These percentile rank scores correspond to how strongly an allele binds a particular peptide relative to a set of random natural peptides. From peptide level rank scores, MHC-I mutation specific binding affinities were assigned according to the best rank score, the minimum allele-specific rank score across all unique 8-11mers for a mutation.

For conserved antigens, we partitioned proteins into their entire set of 8-11mers across the full length of the protein. MHC-I allele peptide pair percentile rank scores were again generated using NetMHCpan-4.1. Several metrics were derived from the percentile rank scores for downstream analyses. For broad and gene-level comparisons, we defined an allele-specific conserved antigen repertoire as the set of 8-11mers presented at or below a given percentile rank (**Fig. 4B-D**, **Supplementary Fig. 10B, 12**). For position-wise presentability (**Fig. 4E-F, Supplementary Fig. 13**) we used the best percentile rank of all overlapping peptides at each position along a protein.

### HLA Population Allele Representations

To compare HLA-B and HLA-C autoimmune (AI) alleles to common non-AI alleles, defined as those alleles with a population frequency ≥ 1% as given by the NMDP (19 common HLA-B alleles, 13 common HLA-C alleles), we established maximum and minimum population allele representations. For driver neoantigens, a maximum population allele representation was assigned to have coverage at each rank equating to the coverage of the best presenting common allele at that rank. Similarly a minimum population allele representation was assigned to have coverage at each rank equating to the coverage of the worst presenting common allele at the rank. For maximum and minimum population allele representations in conserved antigens (CA), the metric of assignment was the fraction of CA peptides bound as opposed to coverage, as was consistent with CA analyses.

### PRS Implementation

We implemented the melanoma PRS developed by Gu *et al.* ^97^ comprising 204 SNPs. For the discovery cohort we were able to extract 190 of the 204 risk SNP genotypes using PLINK ^181^. For the validation cohort, datasets lacking sufficient SNP data for PRS construction were excluded leaving 239 individuals for which 201 of the 204 risk variants were extracted using PLINK2.^182^ For the Melanostrum Consortium (N=3001), all 204 risk SNPs were extracted. Risk SNP MAFs were compared across cohorts to ensure no significant differences (**Supplementary Fig. 7**). For the MVP we were able to extract 202 of the 204 risk variants (rs2025016 and rs6833655 were missing) using PLINK2.^182^ The final PRS for each cohort was generated as a weighted sum across extracted risk SNPs, ensuring SNPs were oriented to the correct allele, in each cohort in accordance with the optimal melanoma risk model by Gu *et al*.^97^

### Multistage Carcinogenesis Model for Melanoma

Dating back to the 1950’s Armitage-Doll model ^183^ of cancer incidence and those created soon after by Knudsen and Moolgavkar ^184^ and others, multistage models of cancer are among the most developed mathematical methods for defining carcinogenesis and determining timescales of tumor formation in human populations.^185–189^ These models assume evolutionary stages from normal cells to development of clinically detected symptomatic cancers. These stages typically include intermediate premalignant and preclinical malignant stages that represent field cancerization dynamics of stochastically growing and shrinking clonal populations in a tissue. These models can be described mathematically as stochastic multi-type branching processes with probabilities of events occurring with certain rates (**Supplementary Fig. 9A**). By calculating an age-dependent hazard function for cancer incidence using solutions to equations from the probability generating functions starting from birth, we can calibrate these models to fit hazard rates derived from cancer incidence registry data such as SEER in the US.^101^ Importantly, this modeling framework provides a link between cell-level dynamics and population-level incidence data so that we can estimate parameters governing clonal growth, dwell times, and mutational “hits” in at-risk individuals.

In previous work we found that the “two-stage” model (2 “hits” for development of a first malignant cell) shown in **Supplementary Fig. 9** is closely approximated by a model that includes an effective malignant transformation rate and a characteristic lag-time or “sojourn” time between malignant transformation and clinical detection (see Luebeck *et al.* 2013 for mathematical details ^100^). Here we created a two-stage model for melanoma incidence that adjusts for birth cohort trends, similar to methods used previously in esophageal squamous cell carcinoma (ESCC).^190^ In this way, our models capture trends for both age and birth cohort (and thus calendar period) to enable robust estimation via Markov Chain Monte Carlo simulation of cell-level parameters for tumor evolution by sex and race/ethnicity (see **Supplementary Fig. 9B** for examples of model fits). We obtained estimates and 95% confidence intervals in the main text for tumor sojourn times in men and women via MCMC posterior estimates for the lag-time parameter. Chains were run for 100,000 cycles with a 4000 cycle burn-in and checked for convergence. All code for hazard function calculation and parameter estimation was written in Fortran. The ICD-O-3 codes used for extraction of SEER data melanoma, all races combined, from SEER*Stat include: 8720/3, 8721/3, 8722/3, 8723/3, 8726/3, 8727/3, 8728/3, 8730/3, 8740/3, 8741/3, 8742/3, 8743/3, 8744/3, 8745/3, 8746/3, 8761/3, 8770/3, 8771/3, 8772/3, 8773/3, 8774/3, 8780/3, 8790/3.

### Statistical Analyses

All box plot statistical tests comparing age of diagnosis effects between groups were assessed using the default Mann-Whitney U statistical test. Leave-one-out analysis was conducted by narrowing the AI allele set into all 7 unique sets of 6 AI alleles and stratifying individuals accordingly. Performing leave-one-out analysis by dropping all carriers of each allele yielded similar results, with AI status significantly associated with a later age of diagnosis in each holdout set. T-tests were used to compare PRS distributions across AI-allele status in both discovery and validation cohorts. Fisher’s exact tests were used to evaluate associations between: 1) MHC-I AI allele carrier status and ICPI response status 2) Major and minor AI SNP alleles and ICPI response status 3) HLA-proximal PRS SNPs and MHC-I AI allele carrier status. These statistical tests were all implemented via the default scipy.stats Python package. Regression analyses were modeled using ordinary least squares linear models through the statsmodels.formula.api Python package.^191^ Effect size for AI alleles and PRS SNPs in the TCGA were calculated using Cliff’s D. Effect size for AI alleles and PRS SNPs in the case-control MVP were reported as odds ratios calculated from logistic regression through the statsmodels.api Python package. All multiple hypothesis testing correction utilized the Benjamini-Hochberg procedure, and was implemented by means of the statsmodels.stats.multitest package in Python.

## Notes

### Competing Interest Statement

The authors have declared no competing interest.

### Summary of Updates

This updated manuscript contains additional analyses - including MHC-I autoimmune allele carrier effect in a large case-control cohort - and co-authors.

