## Supplementary Figures And Tables for "Autoimmune Alleles at the Major Histocompatibility Locus Modify Melanoma Susceptibility"

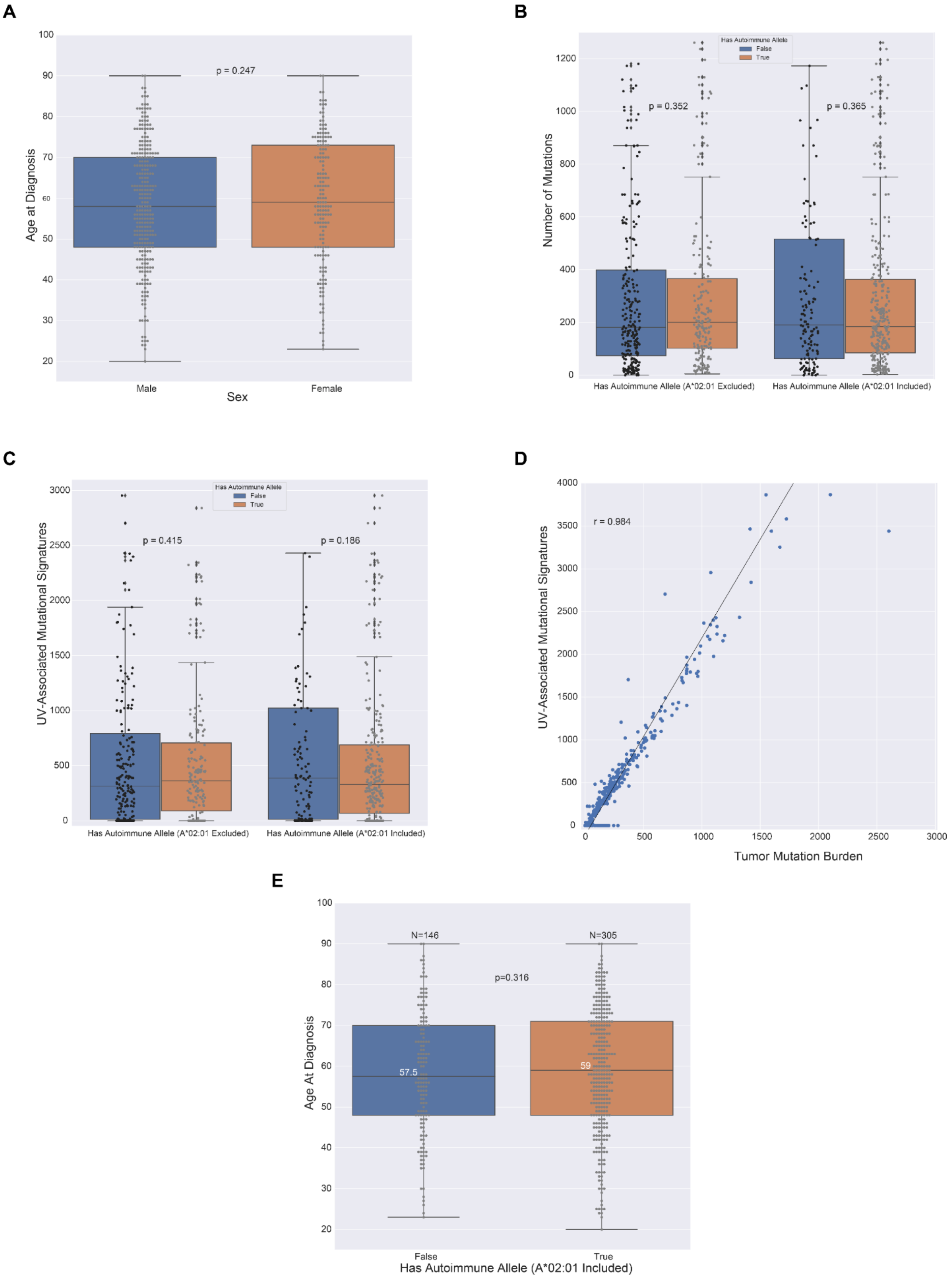
A
p = 0.247
Age at Diagnosis
Male
Female
Sex
B
Has Autoimmune Allele
False
True
p = 0.352
p = 0.365
Number of Mutations
Has Autoimmune Allele (A*02:01 Excluded)
Has Autoimmune Allele (A*02:01 Included)
C
Has Autoimmune Allele
False
True
p = 0.415
p = 0.186
UV-Associated Mutational Signatures
Has Autoimmune Allele (A*02:01 Excluded)
Has Autoimmune Allele (A*02:01 Included)
D
r = 0.984
UV-Associated Mutational Signatures
Tumor Mutation Burden
E
N=146
N=305
p=0.316
57.5
59
Age At Diagnosis
False
True
Has Autoimmune Allele (A*02:01 Included)

**Supplementary Figure 1**: Covariates and associations with melanoma age of diagnosis in the TCGA. A) Boxplots comparing sex and melanoma age of diagnosis. B) Boxplots comparing the number of mutations across those with and without MHC-I linked autoimmune alleles C) Boxplots comparing the number UV-signature associated mutations across those with and without MHC-I linked autoimmune alleles D) Correlation between total mutation burden and UV-signature associated mutations. E) Effect of MHC-I autoimmune alleles on age at diagnosis in melanoma including HLA-A*02:01.

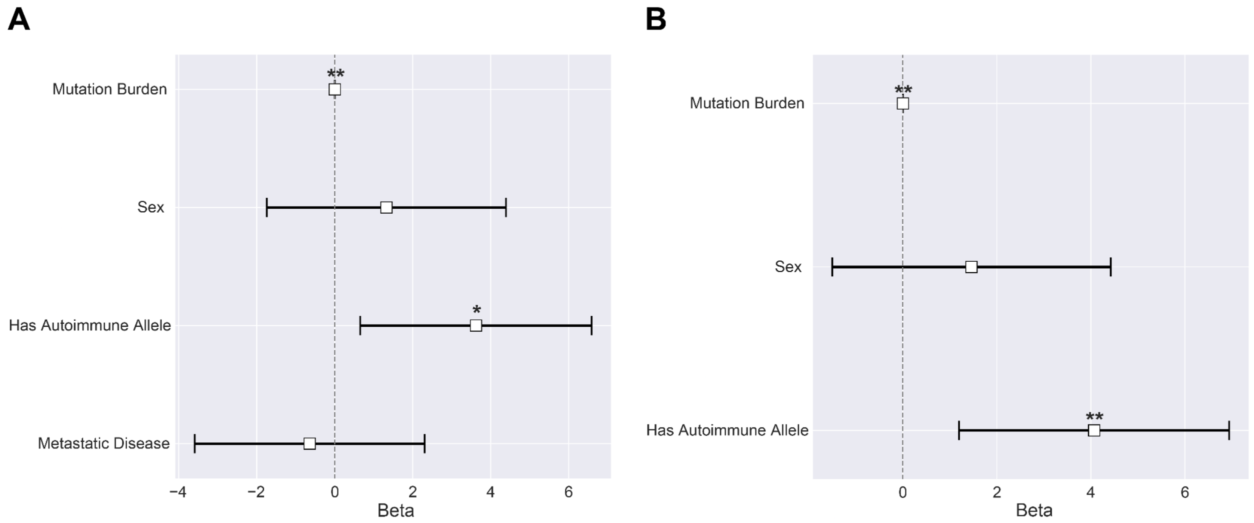


**Supplementary Figure 2:** Regression modeling of AI allele effects on melanoma age of diagnosis in the TCGA. A) Having at least one MHC-I linked autoimmune allele is significantly associated with 3.62 delayed years to melanoma diagnosis after controlling for primary vs. metastatic disease, sex, and mutation burden ($N = 416$, $p_{autoimmune} = 0.015$). Those with an AJCC pathologic tumor stage of II or below were assigned a primary disease label, while those with a tumor stage of III or higher were assigned a metastatic disease label. B) Having at least one MHC-I linked autoimmune allele is significantly associated with 4.07 delayed years to melanoma diagnosis after controlling for sex and mutation burden ($N = 451$, $p_{autoimmune} = 0.005$).

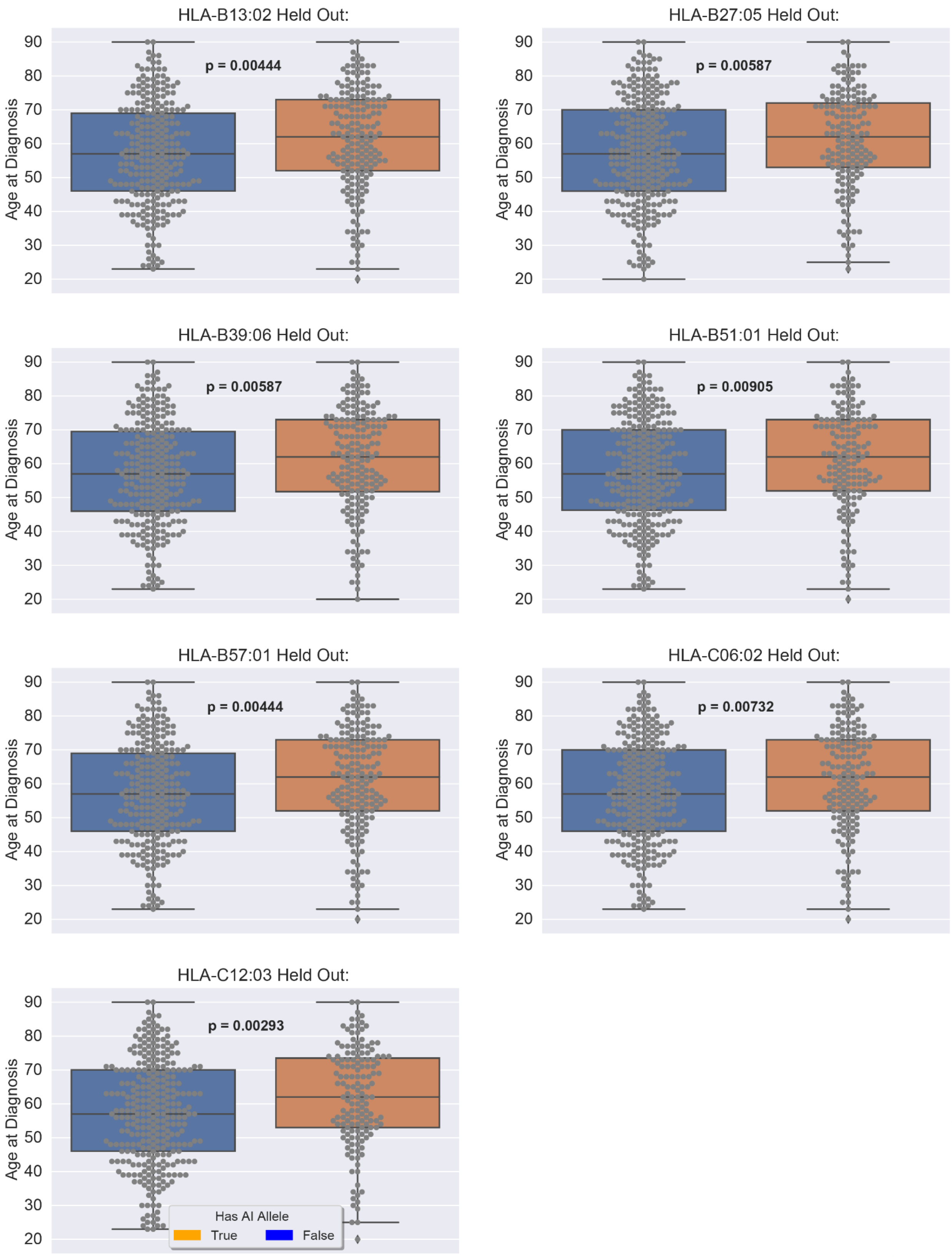
HLA-B13:02 Held Out:
p = 0.00444
HLA-B27:05 Held Out:
p = 0.00587
HLA-B39:06 Held Out:
p = 0.00587
HLA-B51:01 Held Out:
p = 0.00905
HLA-B57:01 Held Out:
p = 0.00444
HLA-C06:02 Held Out:
p = 0.00732
HLA-C12:03 Held Out:
p = 0.00293
Age at Diagnosis
20
30
40
50
60
70
80
90
Has AI Allele
True
False

**Supplementary Figure 3**: Effect of omitting each of the 7 individual AI alleles from carrier status in TCGA. For each AI allele, individuals carrying only the excluded allele were assigned to non-AI allele carrier status.

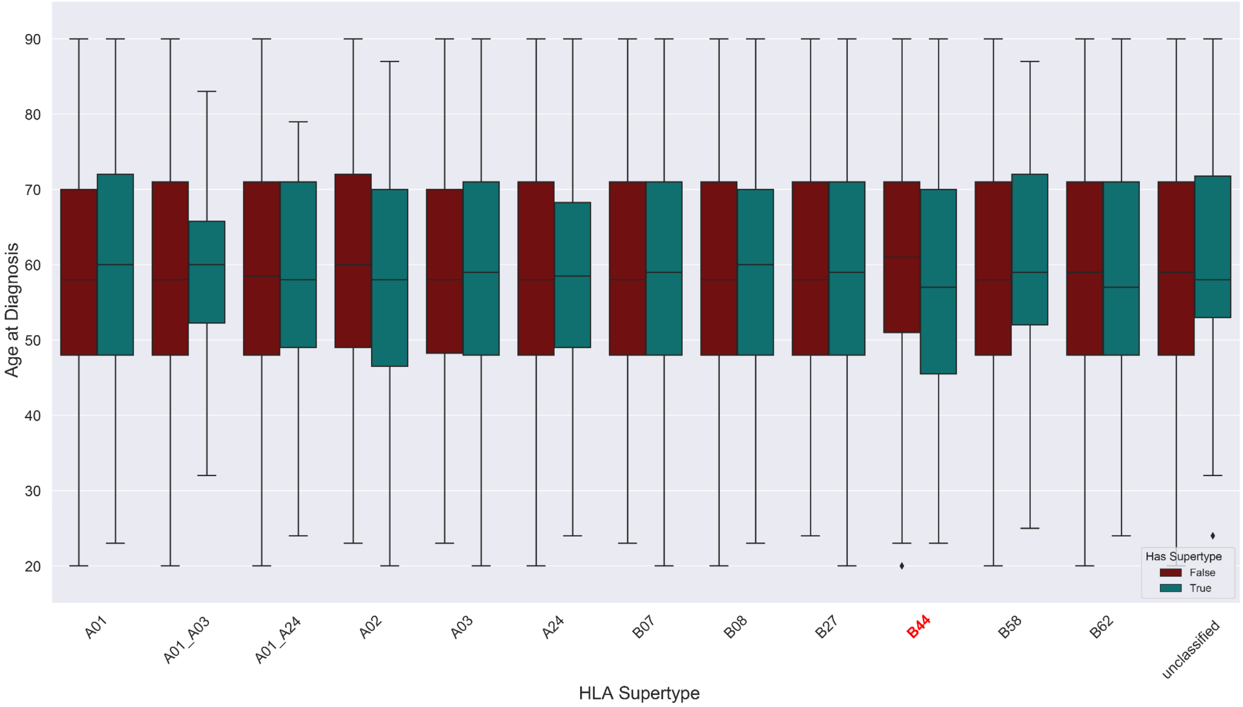


**Supplementary Figure 4**: Effect of HLA supertype on age at diagnosis in the TCGA. No significant age differences were observed across any supertype, though those with the B44 supertype (red) trended towards an earlier age of diagnosis (p = 0.199, median earlier age of diagnosis difference = 4 years).

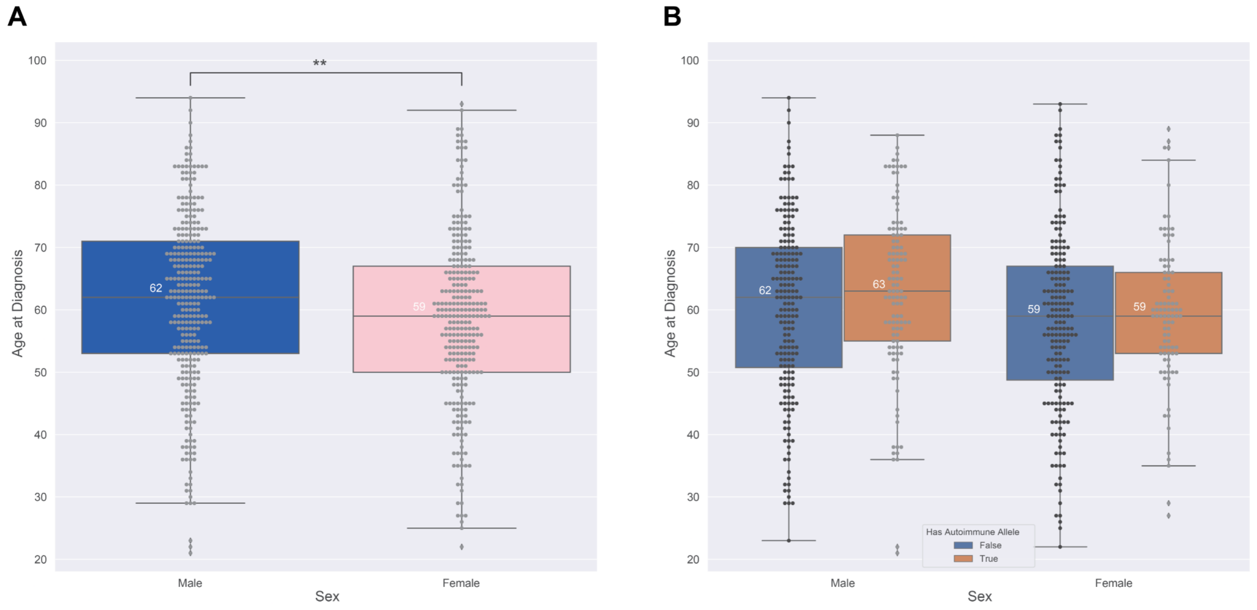


**Supplementary Figure 5**: A) Females in the validation cohort showed a significant earlier age of diagnosis relative to males (p = 0.0033, median difference = 3 years) B) Direction of AI status effect in males is consistent with later age of diagnosis findings ($p_{male}$ = 0.055, $p_{female}$= 0.236).

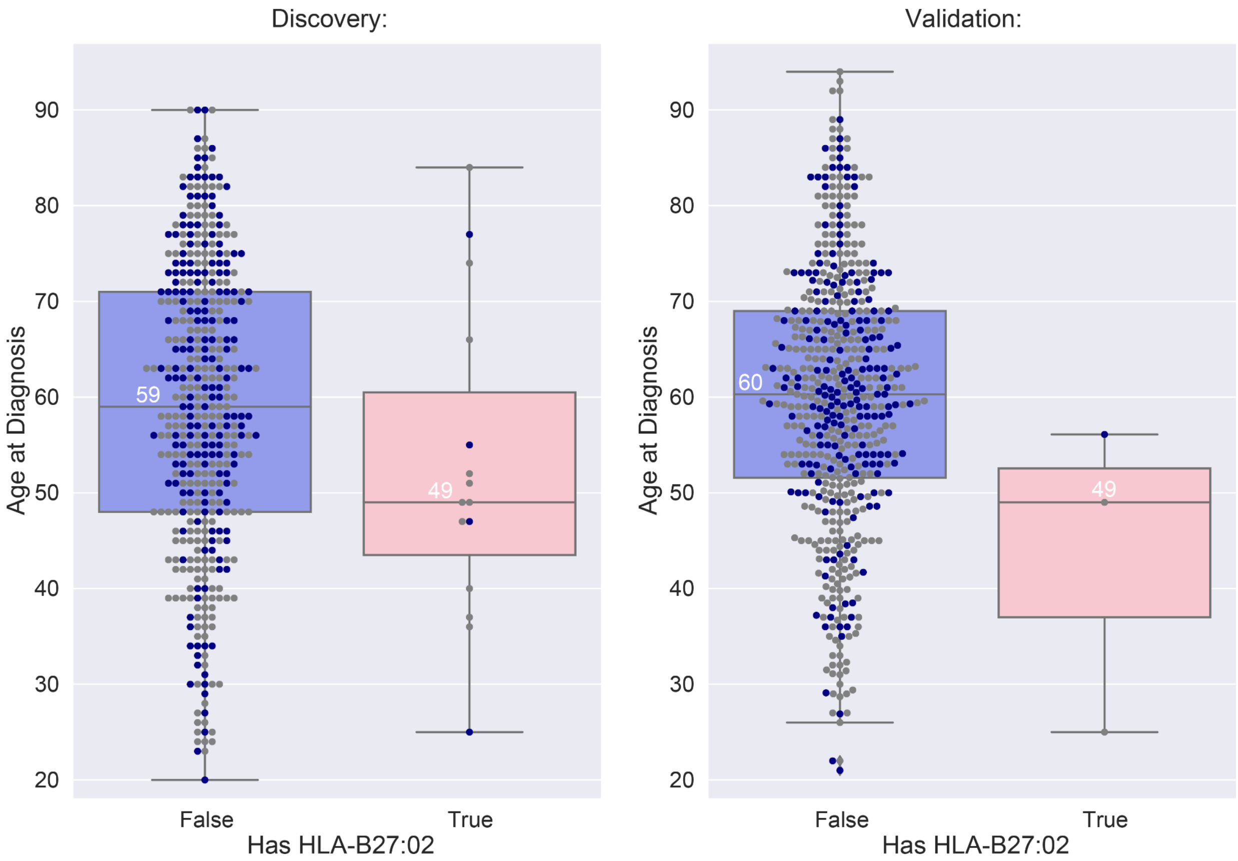


**Supplementary Figure 6**: HLA-B27:02 was associated with an earlier age of diagnosis across both discovery and validation cohorts. Blue points correspond to MHC-I AI allele carriers, while grey points correspond to those without any MHC-I AI alleles.

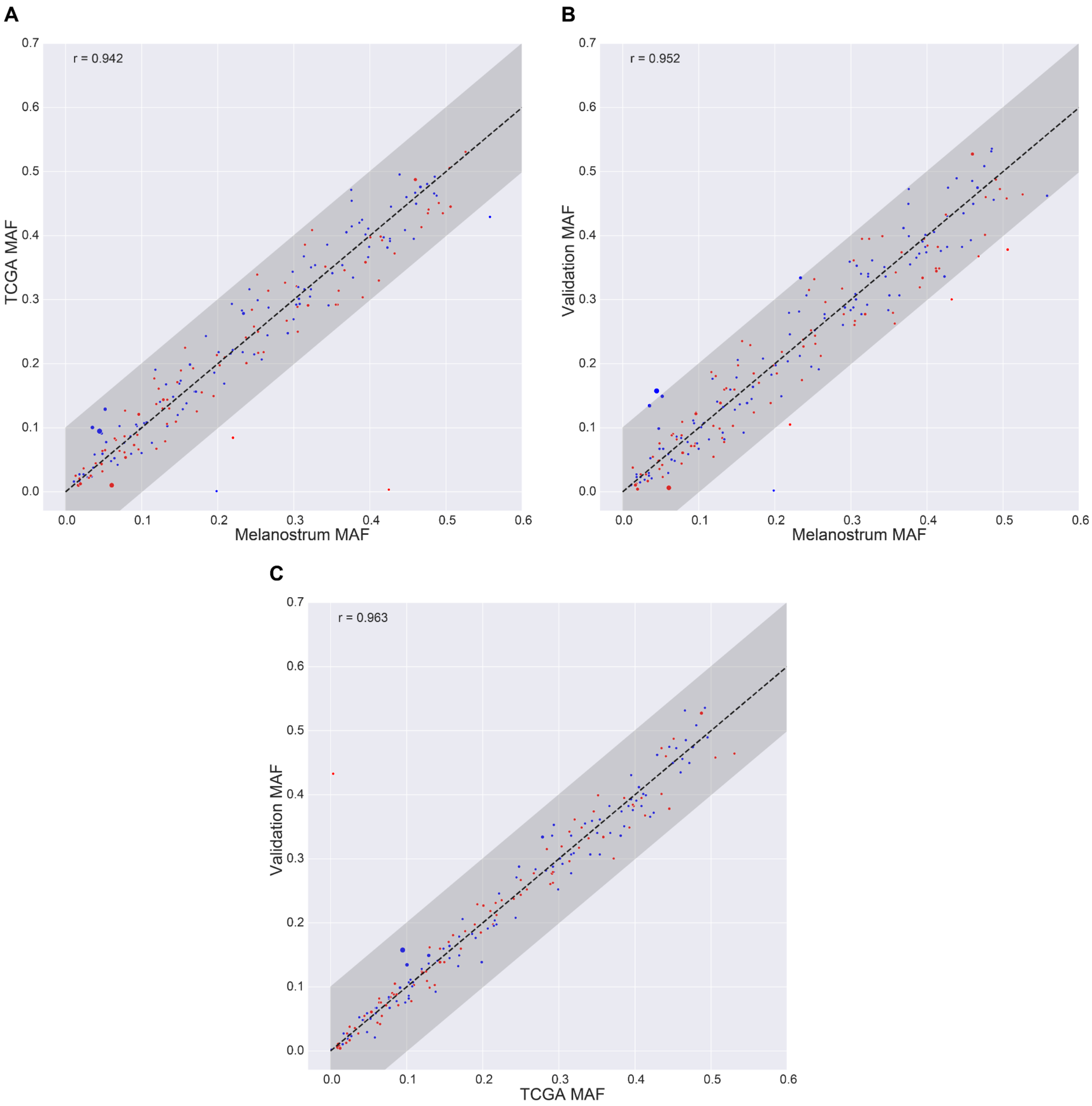


**Supplementary Figure 7**: PRS SNP minor allele frequencies (MAFs) relationship between A) Discovery and Melanostrum (Pearson R = 0.942), B) Validation and Melanostrum (Pearson R = 0.952), and C) Validation and Discovery (Pearson R = 0.963). Red points are PRS SNPs with a protective effect in melanoma (negative PRS weight), while blue points are PRS SNPs with a predisposing melanoma effect (positive PRS weight). Points are sized according to the magnitude of their weight in the PRS. SNP effects ranged in PRS weight from -0.4 to 0.49.

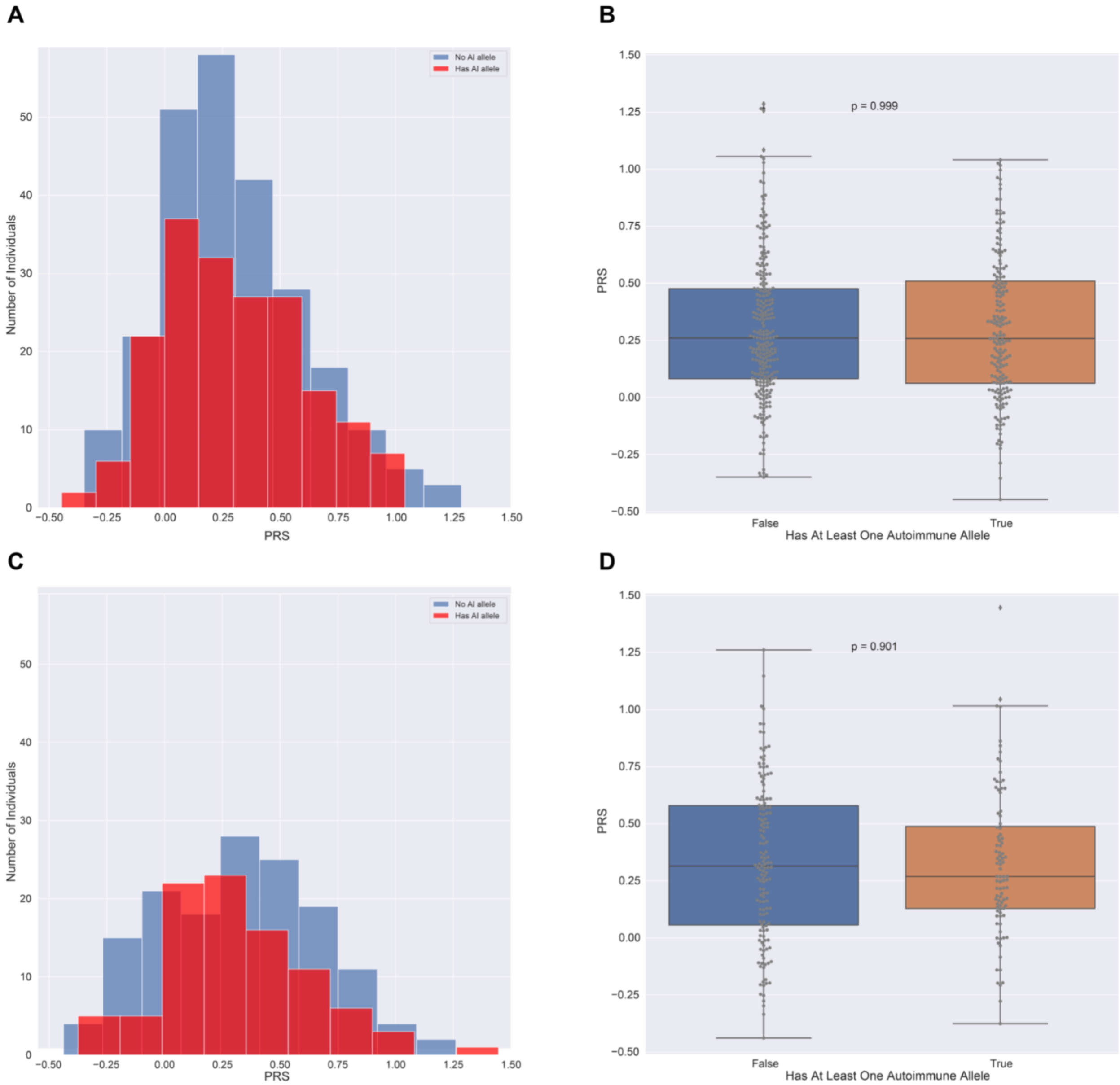
A
B
C
D
No AI allele
Has AI allele
Number of Individuals
PRS
p = 0.999
p = 0.901
False
True
Has At Least One Autoimmune Allele

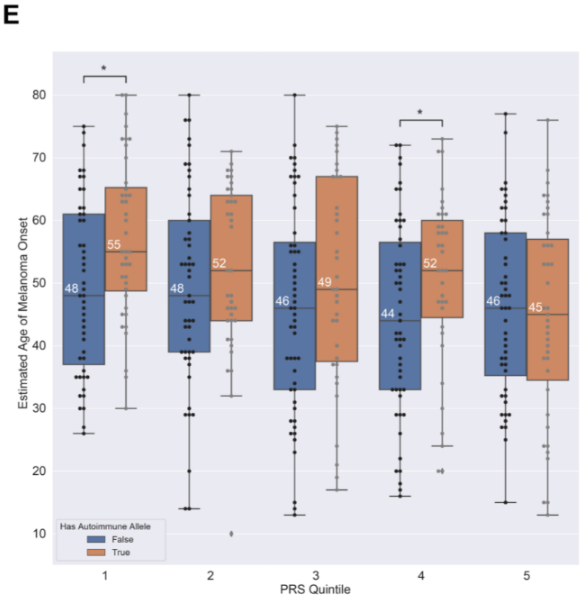
E
Estimated Age of Melanoma Onset
PRS Quintile
Has Autoimmune Allele
False
True

**Supplementary Figure 8**: PRS and MHC-I autoimmune alleles exhibit independent effects with age. A) *TCGA*: Histogram of PRS stratified across individuals with and without MHC-I autoimmune alleles. B) *TCGA*: Boxplots stratifying PRS across individuals with and without MHC-I autoimmune alleles ($p = 0.999$, T-test). C) *Validation*: Histogram of PRS stratified across individuals with and without MHC-I autoimmune alleles. D) *Validation*: Boxplots stratifying PRS across individuals with and without MHC-I autoimmune alleles ($p = 0.901$, T-test). E) *TCGA:* Boxplots of estimated age of melanoma onset across PRS quintiles. Autoimmune carrier status exhibited significant later predicted ages of onset for the lowest (PRS Quintile 1; $p = 0.005$) and second-highest risk quintiles (PRS Quintile 4; $p = 0.034$).

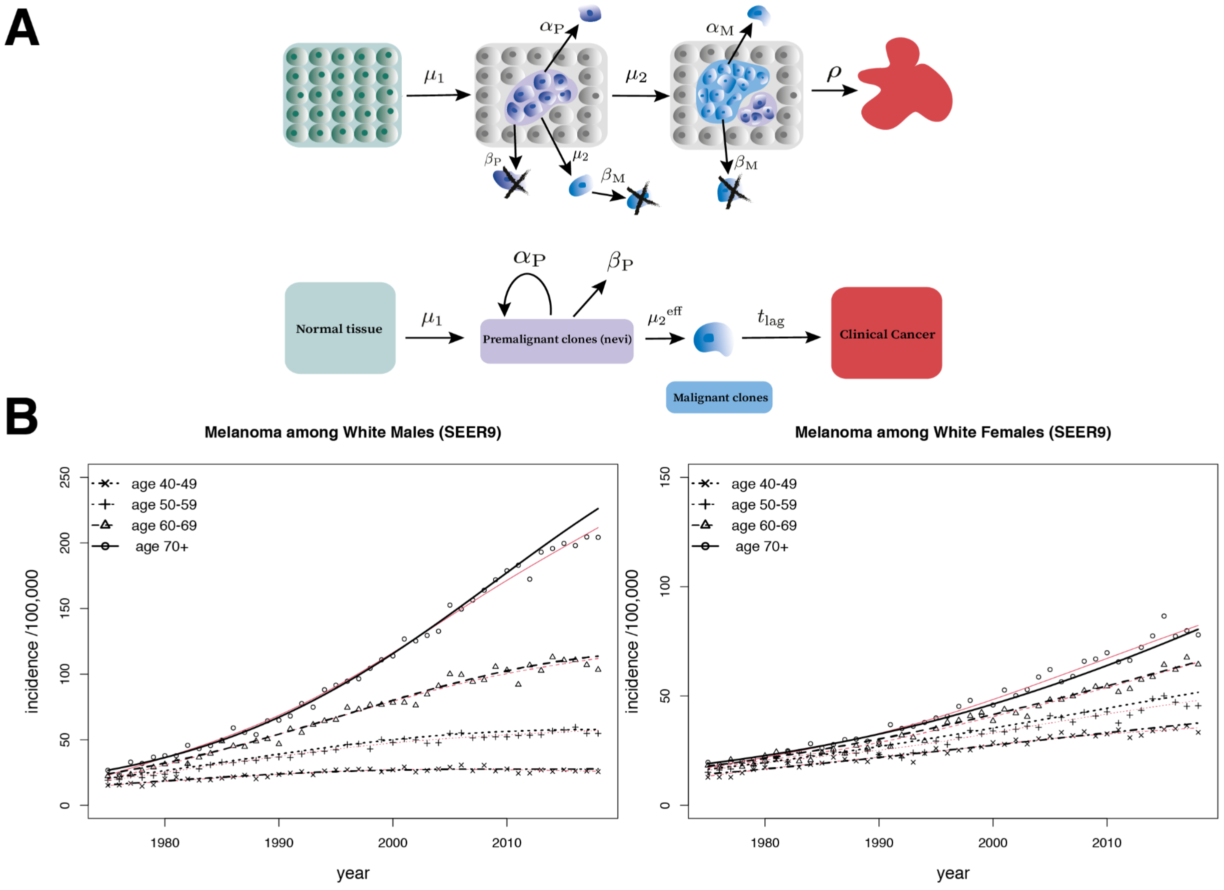


**Supplementary Figure 9**: A) *Upper row*: Multistage clonal expansion (MSCE) model - illustration of a stochastic realization of the multi-type branching process. In the two-stage MSCE model, the first 'hit' occurs when normal cells (green) undergo asymmetric division (due to mutation, for example) and create one premalignant daughter cell with rate $\mu_1$ and one normal cell. Premalignant cells can then clonally expand with rates $\alpha_P$ for division and $\beta_P$ for cell death/differentiation (purple clones, e.g., dysplastic nevi). The second 'hit' occurs at rate $\mu_2$ per cell per year wherein a premalignant cell creates a malignant daughter cell (due to secondary mutation, for example) and 1 premalignant cell. $\alpha_M$ and $\beta_M$ represent cell division and death/differentiation, respectively, for malignant cells (blue). Clinical detection occurs for malignant clones with rate $\rho$. *Bottom row*: Mathematical approximation of the two-stage model includes an effective mutation rate $\mu_2^{eff}$ for transformation of a malignant cell that survives and a tumor sojourn time/ 'lag-time' representing the time between the first persistent malignant cell and the clinically detected cancer. B) Hazard curves from the MSCE models for melanoma incidence/100,000 (black lines) correspond to US SEER data from 1975-2018 (black shapes) in white women (left) and white men (right) separately by age groups, as represented by the different symbols, over calendar years. Calibrated parameters from the two-stage model yield similar average lag-time (tumor sojourn time) for males and females of ~10 years. For comparison, non-parametric fits using 4*th*-order smoothing splines shown in red.

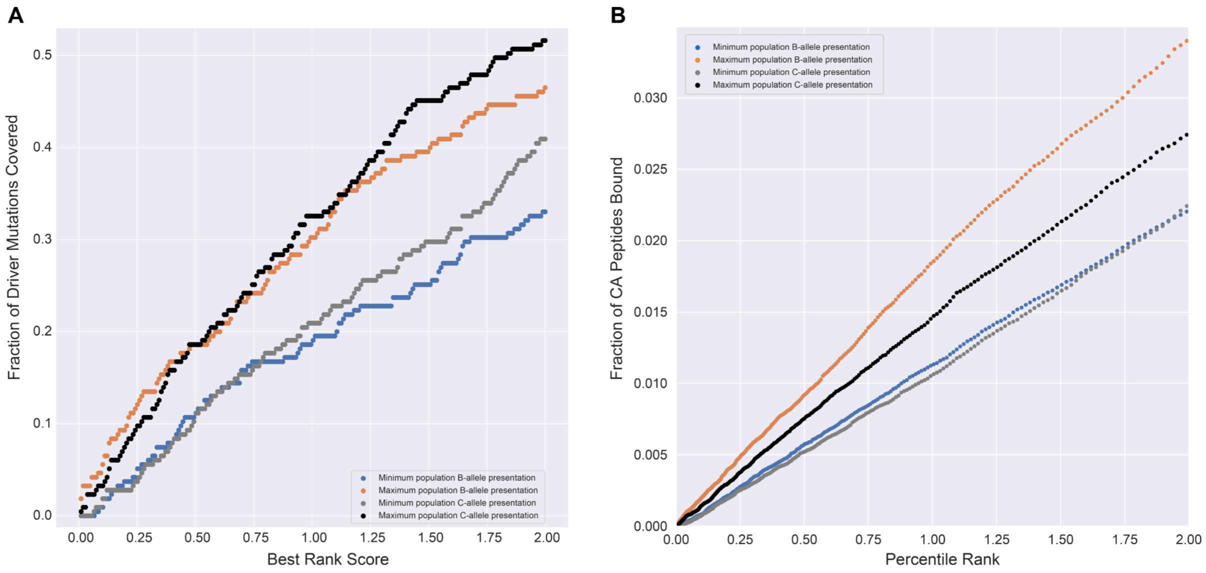


**Supplementary Figure 10:** A) Maximum and minimum HLA-B population allele representations (Methods: *HLA Population Allele Representations*) exhibit greater neopeptide coverage than HLA-C maximum and minimum population allele representations at lower rank scores. As rank scores exceed the classical strong binding threshold (0.5), HLA-C population allele representations generally exhibit greater neopeptide coverage. B) For conserved antigens (CA), the maximum and minimum population allele representation for HLA-B presents a greater fraction of peptides than the maximum and minimum population allele representation for HLA-C. This is most pronounced in the maximum representation, where HLA-B presents a greater fraction of CA peptides compared to HLA-C as percentile rank increases.

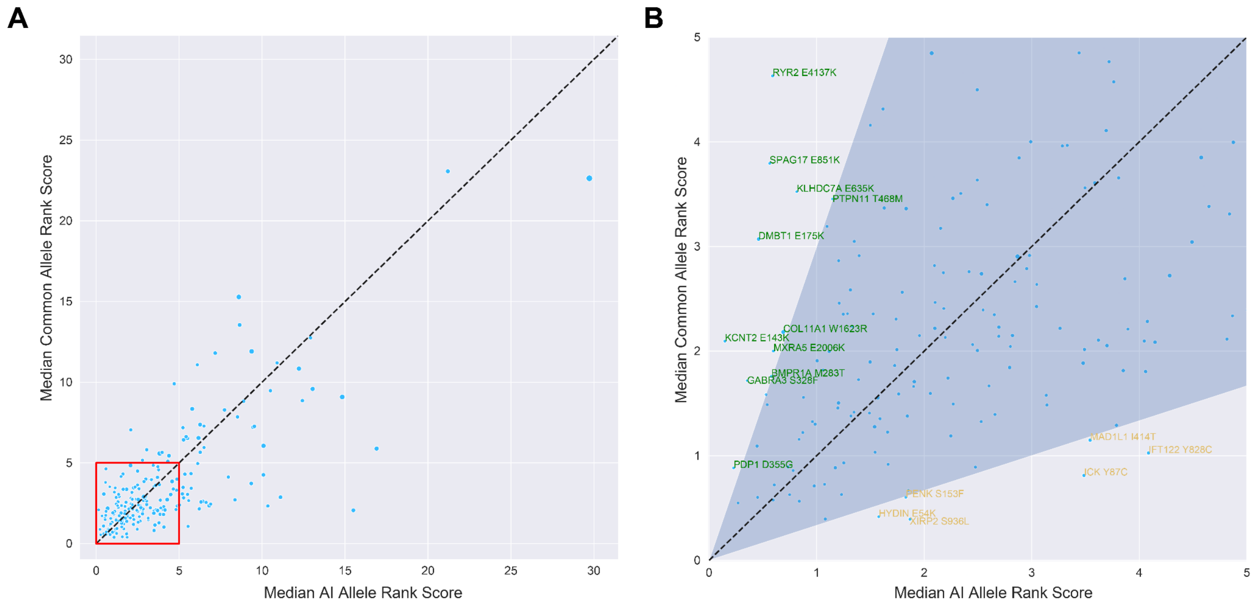


**Supplementary Figure 11**: Predicted median driver mutation rank scores for both common and AI alleles. The varying point size reflects a mutation's standard deviation across all AI alleles, with larger points having larger standard deviations. A) Predicted median rank scores for all driver mutations (N=215) for both common and AI alleles. B) Predicted median rank scores for driver mutations in a presentable range (outlined in red in panel A). Mutations with a 3-fold median rank score discrepancy between AI and common alleles in either direction are labeled, and fall outside the shaded region. Green mutations correspond to mutations that are better presented by the median AI allele, while yellow mutations correspond to mutations that are better presented by the median common allele.

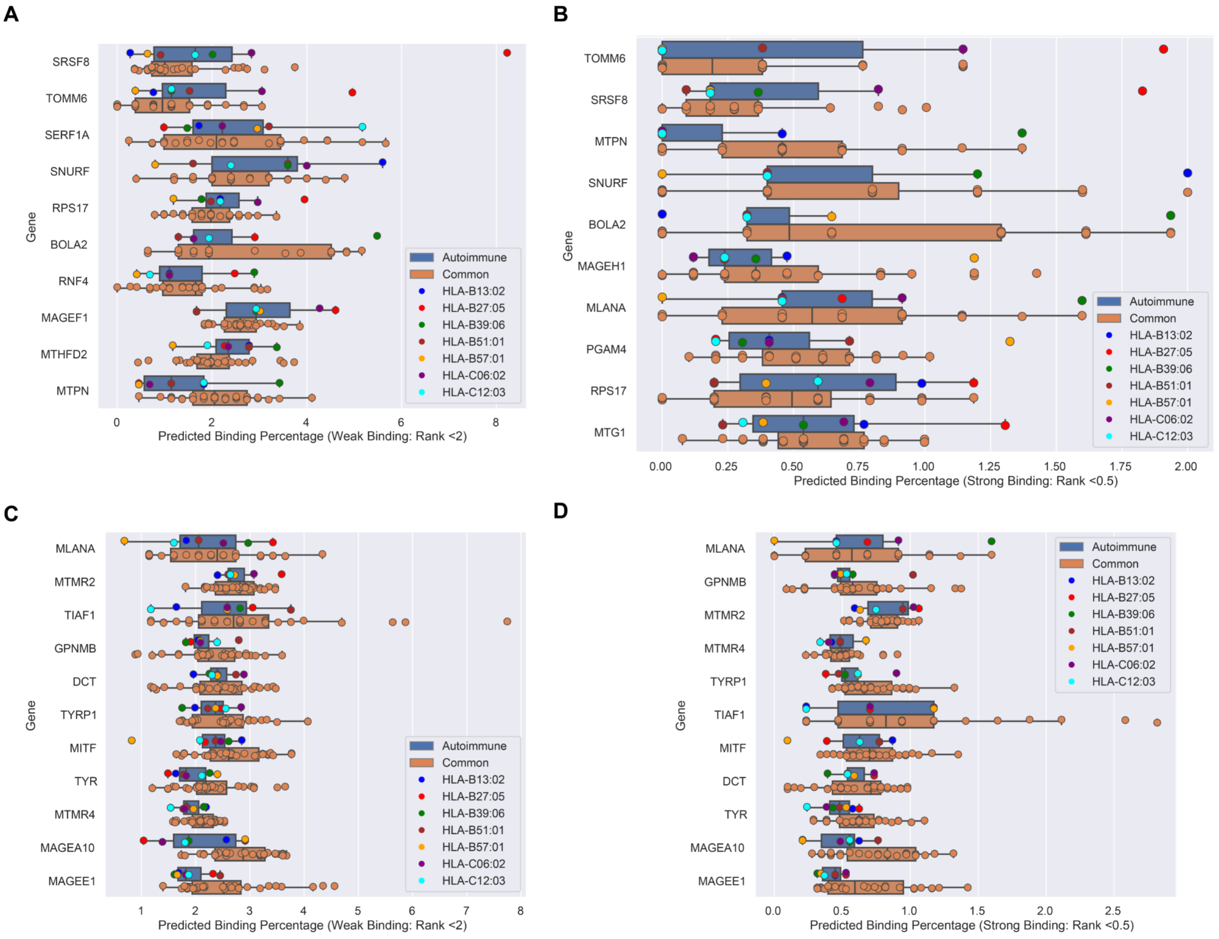


**Supplementary Figure 12**: Distributions of the predicted binding repertoire of conserved antigens between autoimmune and common alleles. A-B) Top 10 conserved antigens by difference in mean % binding (autoimmune - common). Percent binding at the A) 2% rank threshold and B) 0.5% rank threshold. C-D) Differentially expressed conserved antigens sorted as in A-B showing % binding at the C) 2% rank threshold and D) 0.5% rank threshold.

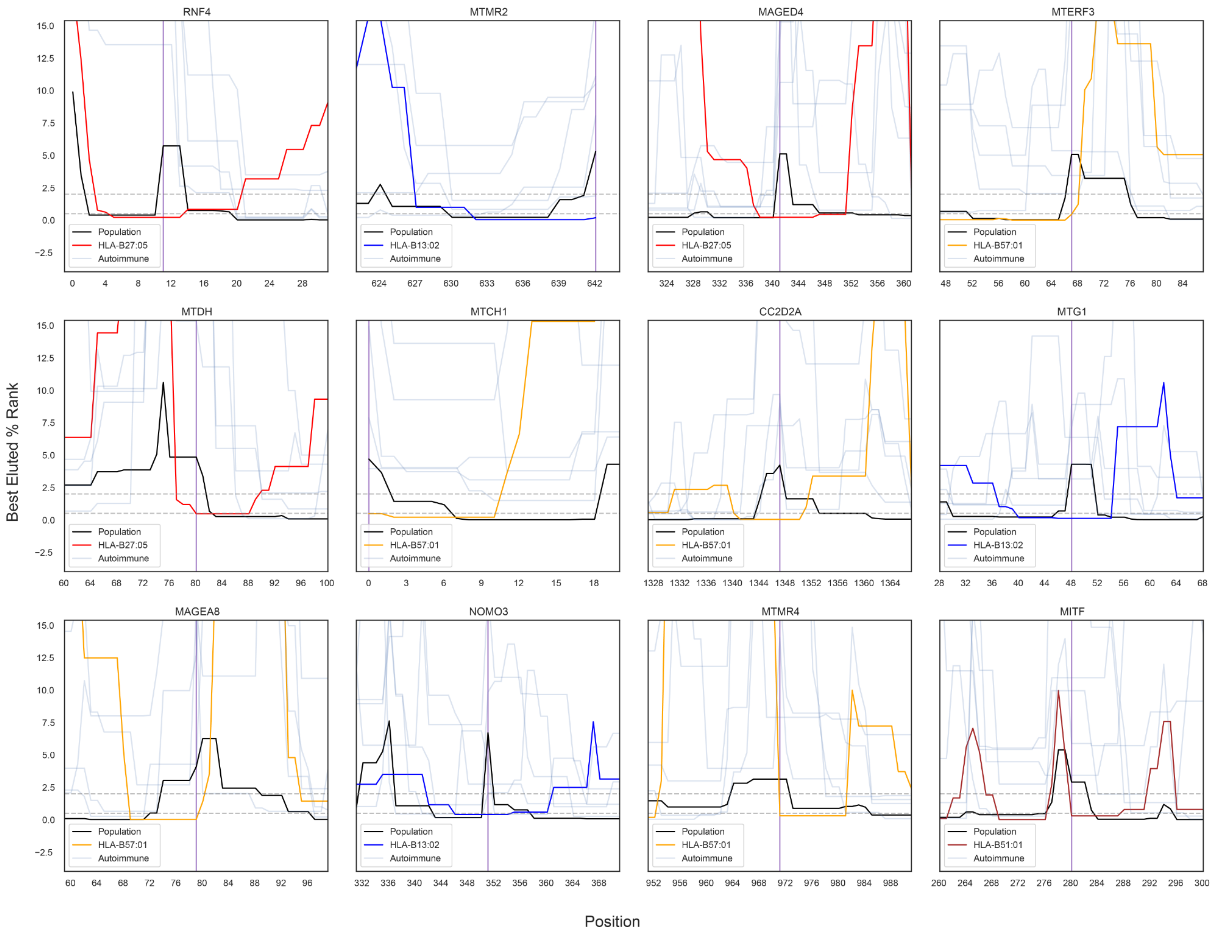


**Supplementary Figure 13**: Greatest position-wise differences in predicted binding to conserved antigens between HLA-B AI and common alleles amongst the top 9 differences (*RNF4, MAGED4, MTERF3, MTDH, MTCH1, CC2D2A, MTG1, MAGEA8, NOMO3*) and DE conserved antigens (*MTMR2, MTMR4, MITF*) not shown in Fig. 4F. Common alleles are shown in orange and AI alleles are shown in transparent blue unless they are predicted to elute at better percentile ranks than common alleles. Vertical purple lines demarcate positions where AI alleles are predicted to elute at better percentile ranks than common alleles and plots show up to +/- 20 aa's of the demarcated positions.

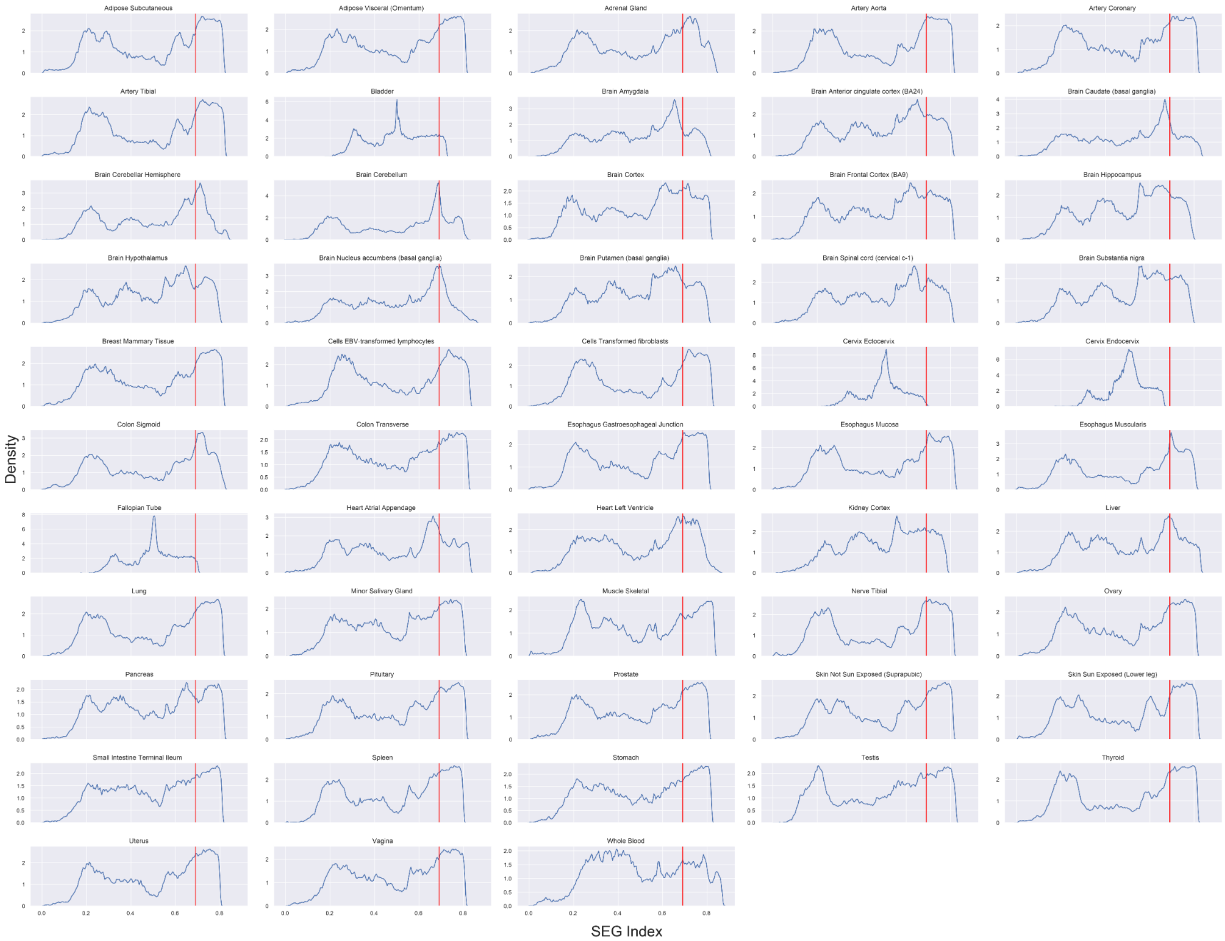


**Supplementary Figure 14**: Distributions of SEG scores shown for all 53 tissues in GTEx. The threshold of 0.69 was selected and is shown in red.

| | Number of Individuals | Average Age (± σ) | Sex (M/F) |
|---|---|---|---|
| TCGA | 451 | 58.66 ± 15.28 | 280/171 (37.9%) |
| Van Allen et al. [92] | 108 | 59 ± 15.96 | 76/32 (29.6%) |
| Hugo et al. [93] | 35 | 61.46 ± 12.93 | 24/11 (31.4%) |
| The Genetic and Transcriptomic Evolution of Melanoma [94] | 39 | 55.49 ± 16.55 | 22/17 (43.6%) |
| Melanoma Exome Sequencing [95,96] | 138 | 66.37 ± 14.01 | 84/54 (39.1%) |
| UKBB | 239 | 56.53 ± 10.47 | 79/160 (66.9%) |

**Supplementary Table 1**: Study specific statistics across both the discovery (TCGA) cohort and validation subsets. Study statistics are reported for individuals ≥ 20 years old, as individuals younger than this were excluded from analyses (See Methods*: Datasets*). Values reported in parentheses correspond to the percentage of females in the particular study.

| | Discovery Cohort (TCGA) Allele Frequencies | Validation Cohort Allele Frequencies | Population Allele Frequency | Autoimmune Disease Association |
|---|---|---|---|---|
| HLA-A*02:01 | 26.27% (201) | 25.31% (240) | 27.6% | Vitiligo [74–77] |
| HLA-B*13:02 | 2.66% (23) | 2.15% (23) | 2.4% | Vitiligo [78,79] |
| HLA-B*27:05 | 3.66% (33) | 4.20% (42) | 3.7% | Psoriasis, Ankylosing Spondylitis [69,70,72,73,154,155] |
| HLA-B*39:06 | 1.00% (9) | 0.63% (7) | 0.64% | Psoriasis, Type 1 Diabetes[69,83–85] |
| HLA-B*51:01 | 4.21% (36) | 3.49% (38) | 4.7% | Psoriasis, Behcet's Disease[82,86] |
| HLA-B*57:01 | 4.21% (38) | 3.22% (35) | 3.6% | Psoriasis [70] |
| HLA-C*06:02 | 9.42% (82) | 7.60% (80) | 9.3% | Vitiligo, Psoriasis [78–81] |
| HLA-C*12:03 | 5.76% (51) | 4.20% (44) | 4.9% | Psoriasis [69,71] |

**Supplementary Table 2**: Autoimmune HLA allele frequency in both discovery (TCGA) and validation cohorts compared against the population distribution as reported by the European Caucasian subset of U.S. National Marrow Donor Program (N = 1,242,890). Values reported in parentheses correspond to the number of individuals in the dataset with the respective allele.

| Conserved Antigen | Class |
| --- | --- |
| SRSF8 | SEGs |
| MTMR3 | SEGs |
| MTRR | SEGs |
| MTDH | SEGs |
| MTFR1L | SEGs |
| MTMR14 | SEGs |
| DLEU1 | SEGs |
| CC2D2A | SEGs |
| MTAP | SEGs |
| MTX2 | SEGs |
| SNURF | SEGs |
| MTFMT | SEGs |
| MTHFSD | SEGs |
| CBWD6 | SEGs |
| PAGR1 | SEGs |
| C2orf15 | SEGs |
| CBWD1 | SEGs |
| MTHFD1L | SEGs |
| MTHFD1 | SEGs |
| SNHG3 | SEGs |
| CYP51A1 | SEGs |
| TOMM6 | SEGs |
| MTCH1 | SEGs |
| PYURF | SEGs |
| DUSP4 | SEGs |
| RPS17 | SEGs |
| MTPN | SEGs |
| MTCH2 | SEGs |
| MTHFD2 | SEGs |
| NOMO2 | SEGs |
| NOMO3 | SEGs |
| PSMA2 | SEGs |
| MTA3 | SEGs |
| TIAF1 | SEGs |
| GPR89B | SEGs |
| DDX47 | SEGs |
| BOLA2B | SEGs |
| PGAM4 | SEGs |
| MTO1 | SEGs |
| MTG1 | SEGs |
| MBD5 | SEGs |

| | |
|---|---|
| MTX1 | SEGs |
| MTMR2 | SEGs |
| MTERFD1 | SEGs |
| MTFR1 | SEGs |
| GTF2H2C | SEGs |
| C1QTNF6 | SEGs |
| MTA2 | SEGs |
| MTMR4 | SEGs |
| RPS17L | SEGs |
| SERF1B | SEGs |
| MTERF3 | SEGs |
| PMEL | Canonical |
| TYRP1 | Canonical |
| GPNMB | Canonical |
| DCT | Canonical |
| MLANA | Canonical |
| TYR | Canonical |
| MITF | Canonical |
| MAGEA1 | MAGE |
| MAGEA10 | MAGE |
| MAGEA11 | MAGE |
| MAGEA12 | MAGE |
| MAGEA2 | MAGE |
| MAGEA3 | MAGE |
| MAGEA4 | MAGE |
| MAGEA6 | MAGE |
| MAGEA8 | MAGE |
| MAGEA9 | MAGE |
| MAGEB1 | MAGE |
| MAGEB10 | MAGE |
| MAGEB16 | MAGE |
| MAGEB18 | MAGE |
| MAGEB2 | MAGE |
| MAGEB3 | MAGE |
| MAGEB4 | MAGE |
| MAGEB5 | MAGE |
| MAGEB6 | MAGE |
| MAGEC1 | MAGE |
| MAGEC3 | MAGE |
| MAGED1 | MAGE |
| MAGED2 | MAGE |
| MAGED4 | MAGE |

| | |
|---|---|
| MAGEE1 | MAGE |
| MAGEC2 | MAGE |
| MAGEE2 | MAGE |
| MAGEF1 | MAGE |
| MAGEH1 | MAGE |
| MAGEL2 | MAGE |
| NDN | MAGE |
| NSMCE3 | MAGE |

**Supplementary Table 3**: The set of 91 genes considered as sources of conserved antigens.
